# Long-term maintenance of patient-specific characteristics in tumoroids from six cancer indications in a common base culture media system

**DOI:** 10.1101/2024.06.10.598331

**Authors:** Colin D. Paul, Chris Yankaskas, Pradip Shahi Thakuri, Brittany Balhouse, Shyanne Salen, Amber Bullock, Sylvia Beam, Anthony Chatman, Sybelle Djikeng, Jenny Yang, Garrett Wong, Isha Dey, Spencer Holmes, Abigail Dockey, Lindsay Bailey-Steinitz, Lina Zheng, Weizhong Li, Vivek Chandra, Jakhan Nguyen, Jason Sharp, Erik Willems, Mark Kennedy, Matt Dallas, David Kuninger

**Affiliations:** Thermo Fisher Scientific, Frederick, MD, United States; Thermo Fisher Scientific, Carlsbad, CA, United States; Thermo Fisher Scientific, Bengaluru, Karnataka, India

## Abstract

Tumoroids, also known as cancer organoids, are patient-derived cancer cells grown as 3D, self-organized multicellular structures that maintain key characteristics (e.g., genotype, gene expression levels) of the tumor from which they originated. These models have emerged as valuable tools for studying tumor biology, cytotoxicity, and response of patient-derived cells to cancer therapies. However, the establishment and maintenance of tumoroids has historically been challenging, labor intensive, and highly variable from lab to lab, hindering their widespread use. Here, we characterize the establishment and/or expansion of colorectal, lung, head and neck, breast, pancreas, and endometrial tumoroids using the standardized, serum-free Gibco OncoPro Tumoroid Culture Medium. Newly derived tumoroid lines (*n*=20) were analyzed by targeted genomic profiling and RNA sequencing and were representative of tumor tissue samples. Tumoroid lines were stable for over 250 days in culture and freeze-thaw competent. Previously established tumoroid lines were also transitioned to OncoPro medium and exhibited, on average, similar growth rates and conserved donor-specific characteristics when compared to original media systems. Additionally, OncoPro medium was compatible with both embedded culture in extracellular matrix and growth in a suspension format for facile culture and scale up. An example application of these models for assessing the cytotoxicity of a natural killer cell line and primary natural killer cells over time and at various doses demonstrated the compatibility of these models with assays used in compound and cell therapy development. We anticipate that the standardization and versatility of this approach will have important benefits for basic cancer research, drug discovery, and personalized medicine and help make tumoroid models more accessible to the cancer research community.

## INTRODUCTION

A common challenge facing cancer researchers is the need for in vitro cancer models representative of the mutational patterns and gene expression levels observed in patient populations. Indeed, differences in mutational burden and gene expression between patient tumors and immortalized cancer cell lines^1–5^ may contribute to drug development bottlenecks in which agents effective in vitro do not translate to complex and multifactorial clinical settings. As a result, there has been increasing interest in utilizing more physiologically relevant cancer models in pre-clinical research, for a more efficient treatment development pipeline^6^ and to reduce animal testing^7^. Tumoroids, also known as cancer organoids, are patient-derived cancer cells that self-organize into three-dimensional (3D) structures in culture and preserve key phenotypic and genotypic features of patient cancers. Our own work comparing two-dimensional (2D) colorectal cancer cell lines with randomly selected primary colorectal cancer donor samples and non-matched colorectal tumoroids demonstrated that the 2D lines harbored higher numbers of clinically relevant mutations and had higher tumor mutational burdens compared to tumors and tumoroid lines (**Supplementary Fig. S1**). In addition, quantification of gene expression levels followed by hierarchical clustering revealed that 2D lines clustered separately from the primary tumors, which instead clustered together with tumoroids (**Supplementary Fig. S1**). These observations emphasize that 2D lines are, in many cases, hypermutated and are not representative of patient cancer populations compared to tumoroid models. Therefore, tumoroids hold great promise in both pre-clinical and translational research. Specifically, these physiologically-relevant models could reveal aspects of tumor biology that are not recapitulated by immortalized cancer cell lines and could help increase the efficiency of treatment discovery pipelines^8^, and, potentially, identify effective patient treatments through functional precision medicine approaches^9^.

While the value of tumoroid culture in cancer research is apparent, the complexity of the approach has limited its adoption. Many methods for tumoroid culture rely on complicated 3D culture handling and medium production protocols^10^, lack of standardization amongst bespoke media recipes (“homebrew” media systems), reliance on conditioned medium^11–13^, and growth competition with benign cell types^14–16^. In this study, we demonstrate the capability of the serum-free, conditioned medium-free Gibco™ OncoPro™ Tumoroid Culture Medium (referred to throughout as “OncoPro medium”) to support derivation and long-term culture of colorectal, lung, breast, and endometrial tumoroid models, wherein the patient-specific characteristics identified from primary tumor tissue are recapitulated throughout the culture period. The system relies on a conserved base medium, with the addition of 1-2 niche factors for some indications to support growth. Tumoroid expansion in the system is compatible with 3D encapsulation in basement membrane extract (BME) domes but also amenable to a suspension culture approach where the cells float freely in the presence of diluted BME to provide extracellular matrix (ECM) cues. Suspension tumoroid culture reduces BME consumption and simplifies handling to facilitate scale up and use in high-throughput applications. We show the compatibility of this novel media system and suspension culture workflow with the expansion of colorectal, lung, head and neck, breast, and pancreas tumoroid lines that had been previously cultured in homebrew media formulations, while maintaining the distinctive mutational and gene expression profiles of each cell model. Finally, to illustrate the utility of tumoroid models in preclinical cell therapy research, we developed a tumoroid line that stably expresses green fluorescent protein (GFP) and used these tumoroids to interrogate the killing potential of primary natural killer (NK) cells and an immortalized NK cell line (NK-92). Together, our results illustrate how OncoPro medium enables the use of tumoroid models in cancer research by providing a standardized formulation that is broadly effective in helping maintain donor-specific characteristics during in vitro cell culture.

## RESULTS

### Derivation of tumoroid lines from primary cancer samples across multiple indications using a conserved base medium formulation

To demonstrate the effectiveness of our approach for tumoroid derivation, we tested whether the OncoPro medium formulation supported generation of novel patient-derived cultures from viable cancer cells. Dissociated cells, either from commercially available cryopreserved material or following enzymatic dissociation of fresh primary tumor surgical resection samples, were cultured in OncoPro medium **(Figure 1a).** Samples from the initial material were also collected for subsequent genetic and transcriptomic analysis. The indications represented in the derivation experiments were colorectal, lung, breast, and endometrial cancers. For lung cancer samples, the base OncoPro medium formulation was supplemented with Heat Stable FGF-10 (HS FGF-10), while medium for breast and endometrial cancer tumoroids was supplemented with HS FGF-10 and beta-estradiol (**Supplementary Table S1**). Our derivation workflow typically utilized a hybrid approach combining embedded culture in BME hydrogel domes for the first several passages (typically ∼5), followed by scale up in a suspension culture format wherein 2% (v/v) basement membrane extract was added to OncoPro medium in non-tissue culture treated cell culture dishes and flasks to maintain the self-organized 3D morphologies characteristic of tumoroids.

**Figure 1.**
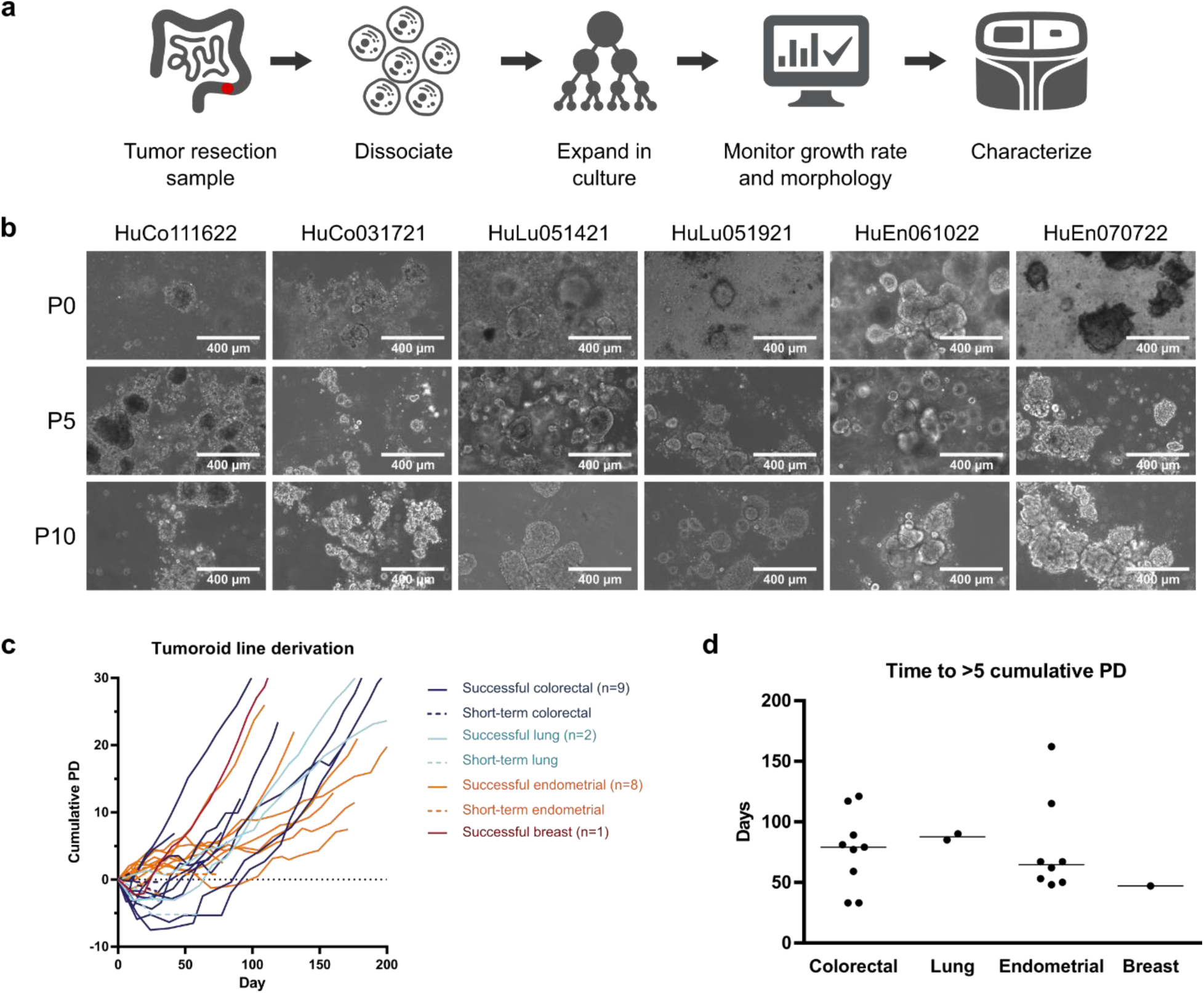
Tumoroid lines can be derived from donor material across several cancer indications. **(a)** Schematic of tumoroid establishment and characterization. Donor tumor material was processed, cultured, and analyzed for fidelity to the original cells sampled at the time of collection by next-generation sequencing. **(b)** Representative images of tumoroid lines at passages (P) 0, 5, and 10. P0 images were acquired after 1–2 weeks of culture. P0 and P5 images represent a mix of embedded and suspension samples. All P10 images were suspension cultures. HuCo = human colorectal cancer; HuLu = human lung cancer; HuEn = human endometrial cancer. Scale bar = 400 μm. **(c)** Cumulative population doublings (PD) over time during attempted derivation of colorectal, lung, endometrial, and breast tumoroid lines. Dashed lines indicate short-term cultures that formed tumoroids initially but failed to expand in vitro over the long term. **(d)** Time (in days) for successfully established tumoroid lines to reach 5 PD based on initial cell seeding number for various indications. Each point represents one donor, and each line is plotted at the median.

Greater than 85% of samples initially formed 3D tumoroid structures and exhibited a range of donor-specific morphologies (**Figure 1b**), which were retained for the duration of culture in either embedded or suspension formats. Tumoroids were subcultured by dissociating multicellular 3D structures when they reached an average diameter of 100-300 µm and seeding a mixture of single cells and small cell clusters (**Supplementary Fig. S2**); samples that did not grow as large were passaged after a period of 14 days. Occasionally, overall cell number would decline during the first few passages in culture (**Figure 1c**). Approximately half of cultures grew stably in the long term, which we defined as expanding consistently for more than 5 passages and exceeding 5 cumulative population doublings from the initial number of cells seeded (including any non-cancer cells present in the initial sample; **Figure 1c**). These stringent cutoffs were good predictors of whether a tumoroid culture would continue to grow consistently and were confirmed in 2 colorectal and 2 lung samples, which grew for over 275 days. Select examples of unsuccessful derivation (tumoroids reforming for only a few passages) are also given (**Figure 1c**). Derivation of cultures to these standards typically required 50-120 days (**Figure 1d**). Based on these derivation criteria, 9 colorectal tumoroid lines, 2 lung tumoroid lines, 1 breast tumoroid line, and 8 endometrial tumoroid lines were successfully established and cryopreserved for subsequent use. Donor characteristics for these derived lines are summarized in **Supplementary Table S2**.

### Genomic characteristics of original tumor cells are retained in patient-derived tumoroid cultures

To help ensure that tumoroids preserve key genomic features of donor cancer material, we performed genomic characterization of successfully derived tumoroid cultures after 5-13 passages using the Ion Torrent™ Oncomine™ Comprehensive Assay v3 (OCAv3) for comparison to initial samples. Quantification of the variant allelic frequency (VAF) of single nucleotide variants (SNVs) detected by the assay revealed high concordance between matched tumors and tumoroids (Pearson correlation coefficient *r* ≥ 0.85) for colorectal (8/9), lung (2/2), breast (1/1), and endometrial (7/8) cancers (**Figure 2a, top panel**). SNV profiles were patient-specific, with little cross-correlation across donors (*r* ≤ 0.70 between donors when all compared; see **Supplementary Table S3** and **Methods**). The proportion of single base substitutions was also retained when comparing original tumor samples to corresponding tumoroids (**Figure 2a, middle panel**). Furthermore, unique sets of oncogenic driver mutations present in the original tumor samples were observed in tumoroid culture (**Figure 2a, bottom panel**). On a per-mutation basis, >87% of SNVs were shared between the initial samples and the derived tumoroids for 19/20 samples, suggesting minimal clonal loss or expansion in culture (**Figure 2b**). The *APC* gene is commonly mutated in colorectal cancers^17^; as this gene is not covered by the OCAv3, we verified the presence of APC mutations in select colorectal tumoroids using quantitative polymerase chain reaction (qPCR) genotyping assays (**Supplementary Table S4**). Overall, these results indicate that the tumoroid media system can support samples across broad genomic landscapes from multiple cancer indications.

**Figure 2.**
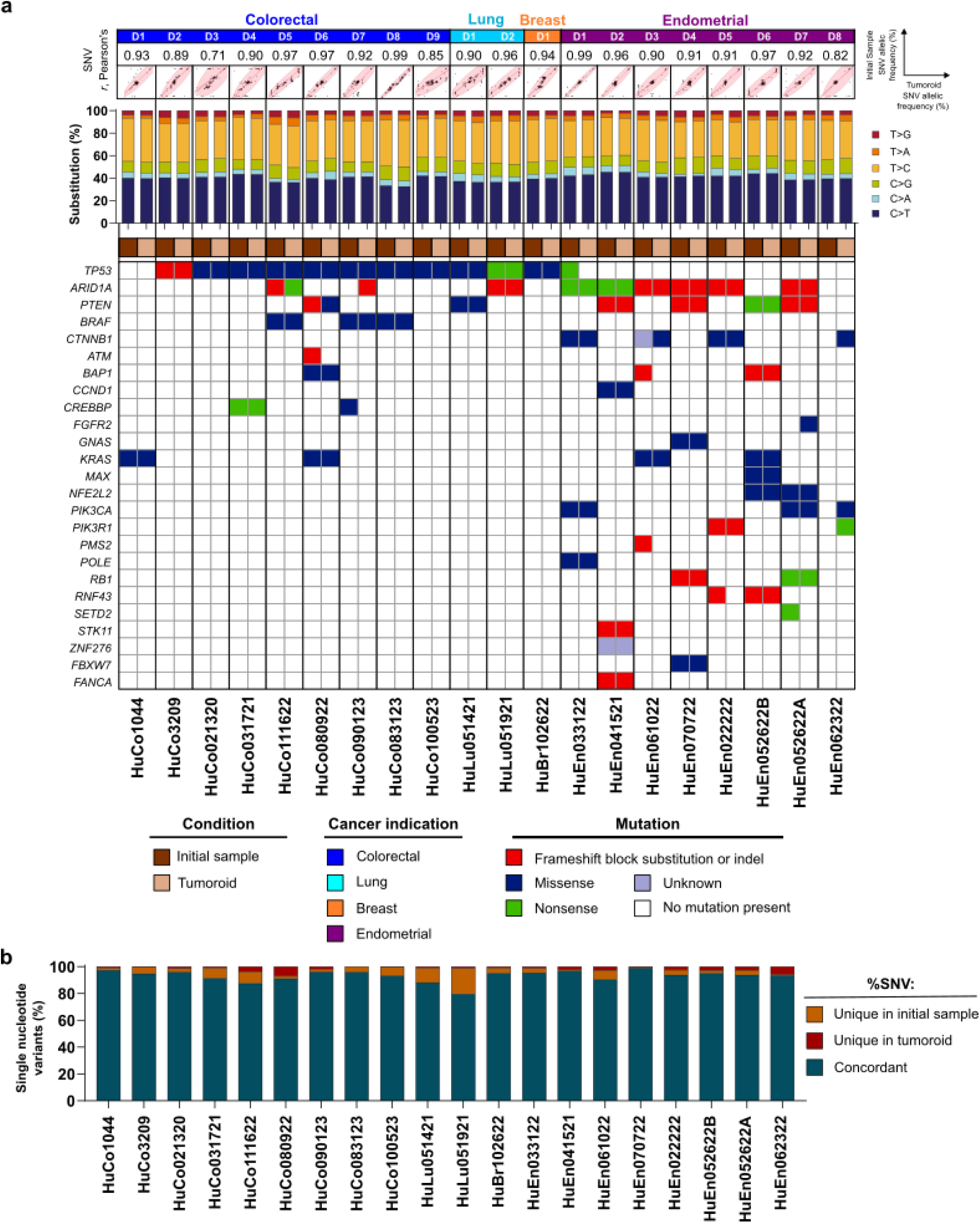
Tumoroids preserve donor-specific mutations from originating material. (**a**) Correlation of the allelic frequency of single nucleotide variants (SNVs) for matched tumor tissue and tumoroid pairs (top), bar plots of single nucleotide base substitutions (middle), and oncogenic single nucleotide variants (bottom) for donor-matched tumor/tumoroid sets for the indicated cancer types. In top panel, each dot represents the variant allelic frequency (VAF) for 1 genetic locus covered by the Oncomine™ Comprehensive Assay v3 (OCAv3). The oncogenic mutations were called using the Oncomine™ Variants filter 5.18 within the Ion Reporter™ software. (**b**) Overlap in SNVs detected between initial donor material and established tumoroid lines (concordant SNVs, dark blue), and SNVs uniquely detected in each sample (orange, red) on a per-donor basis.

### Donor-specific transcriptomic profiles are preserved in tumoroid cultures

To analyze how closely transcriptomic profiles observed in human cancer samples resembled those from donor-matched in vitro tumoroid models, bulk RNA profiling was performed using the Ion Torrent™ Ion Ampliseq™ Transcriptome Human Gene Expression Kit to quantify expression levels of >20,000 human reference sequence genes in tumor and tumoroid pairs. Average Pearson’s correlation values to compare log-transformed, normalized gene read levels between matched tumor and tumoroid pairs of *r*=0.86 (colorectal, n=9), *r*=0.89 (lung, n=2), and *r*=0.87 (endometrial, n=8) were observed, suggesting high transcriptomic similarity (**Figure 3a**). Additionally, the derived breast cancer tumoroid line showed a gene expression level correlation of *r*=0.87 between initial cells and derived tumoroids. Overall, transcriptomic similarity was highest within a given cancer indication compared to cross-indication comparisons (**Figure 3a**).

**Figure 3.**
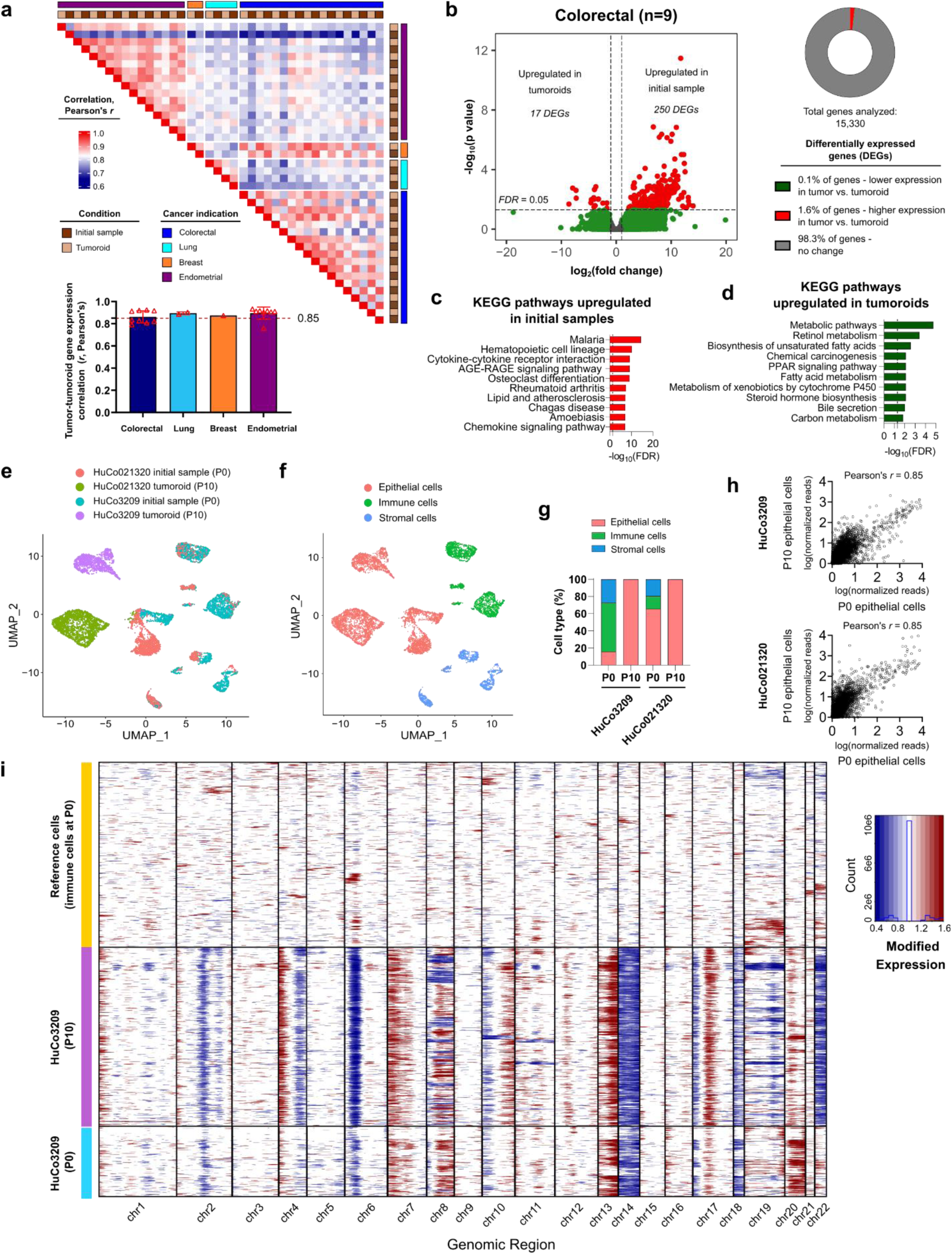
Tumoroids preserve transcriptomic features of original tumors. (**a**) Correlation heatmap of bulk RNA expression levels (17,556 genes) in matched tumor tissue and tumoroids for colorectal, lung, breast, and endometrial cancers. Inset shows average (bar) Pearson’s correlation for matched tumor tissue and tumoroid pairs (dots denote unique donors) across different cancer indications. (**b**) Differential gene expression analysis in matched colorectal tumor tissue and tumoroids (n=9) at fold change>2 and false discovery rate (FDR)<0.05. (**c**) Top 10 significantly enriched Kyoto Encyclopedia of Genes and Genomes (KEGG) pathways of highly expressed genes in initial samples (dotted line indicates FDR=0.05). (**d**) Top 10 significantly enriched KEGG pathways associated with genes highly expressed in colorectal tumoroids (dotted line indicates FDR=0.05). (e) Uniform Manifold Approximation and Projection (UMAP) for dimension reduction visualization of 10,139 single cells from two matched tumor tissue/tumoroid pairs with color coded assignment of samples after single-cell RNA sequencing (scRNA-seq) for colorectal samples HuCo021320 and HuCo3209. (**f**) UMAP visualization with color-coded assignment of cancer cells, stromal cells, and immune cells. (**g**) Proportion of cell types present in initial samples (P0) and matched tumoroids (P10) for two colorectal cancer donors, HuCo021320 and HuCo3209. (**h**) Comparison of normalized, natural log transformed gene expression levels from pseudo-bulked scRNA-seq data of epithelial cells present in initial (P0) and tumoroid (P10) cells for HuCo3209 colorectal cells (17,731 genes) and HuCo021320 colorectal cells (17,536 genes). (**i**) Single cell copy number variation (CNV) plots comparing immune cells, tumor epithelial cells, and tumoroids for HuCo3209 donor. InferCNV was used to estimate CNV in epithelial cells using immune cells as the reference.

Differential gene expression analysis of primary tumors and tumoroids after expansion to establish early cryobanks (passage 3-16, depending on growth rate and initial number of cells seeded) showed minimal changes in gene expression levels associated with tumoroid establishment. Comparison of original tumors and colorectal tumoroids (n=9) showed 267 differentially expressed genes (DEGs) at fold change>2 and false discovery rate (FDR)<0.05, accounting for less than 2% of analyzed genes (**Figure 3b**). Pathway analysis of DEGs highly expressed in tumors but downregulated in tumoroids revealed that the top 10 most enriched Kyoto Encyclopedia of Genes and Genomes (KEGG) pathways were primarily immune cell signaling pathways (**Figure 3c**). Similarly, minimal transcriptome changes of 4.4% and 1.0% were observed when comparing initial tumor cells to established tumoroids in lung cancer (n=2) and endometrial cancer (n=8), respectively, and differences in gene expression were primarily related to immune cell-related signaling pathways (**Supplementary Fig. S3**). Likewise, an unbiased gene expression analysis to identify the top 50 variable genes in our tumor/tumoroid data set followed by hierarchical clustering illustrated a decrease in gene expression levels associated with immune and stromal cells (e.g., *CCL3*, *CCL4*, *COL1A2*, *COL1A3*) when comparing tumors and derived tumoroids (**Supplementary Fig. S3**). This analysis showed retention of expression levels associated with epithelial cancer cells (e.g., *CDH17*, *KRT20*, *CDX1*, *NOX1*, *ESR1*, *SCGB2A1*, *SCGB1D2*, *PAX8*; **Supplementary Fig. S3**). Flow cytometry comparing cells dissociated from a primary tumor to its matched tumoroid culture confirmed the enrichment of EpCAM-positive and CEACAM-positive cells and a reduction in CD45-positive and CD31-positive cells during tumoroid establishment (**Supplementary Fig. S3**). The dropout of immune cell signatures is expected in established tumoroid cultures, as OncoPro medium is intended to culture cancer cells and is not expected to support immune and stromal cells over the long term. Few genes (<0.25%) were differentially upregulated in tumoroids for all three cancer indications, and those genes were primarily related to metabolism signaling pathways (**Figure 3d; Supplementary Fig. S3**).

To directly compare the transcriptome of cancer epithelial cells from tumors and tumoroids, we performed single-cell RNA sequencing (scRNA-seq) of two dissociated tumor cell samples, HuCo3209 and HuCo021320, and matched tumoroid banks at passage 10 (**Figure 3e,f**). Cells from the original tumor material formed distinct clusters of epithelial, immune, and stromal cells (as defined by expression of canonical marker genes; **Supplementary Fig. S4**) that were often nearby or overlapping between the two donors (**Figure 3e,f**), though the proportion of cell types varied by donor (**Figure 3g**). Established tumoroid cells were nearly all epithelial in nature and clustered most closely with the epithelial cells found in the initial sample (**Figure 3e-g**). To directly compare gene expression levels in tumoroids with those in the epithelial cells present initially, scRNA-seq data from the epithelial cell subpopulations in initial samples (P0) and in tumoroids (P10) were pseudo-bulked and compared. Log-transformed normalized read counts between these populations were highly correlated for both HuCo3209 and HuCo021320 samples (*r*=0.85; **Figure 3h**).

Additionally, we investigated if colorectal tumoroids represent the original tumor at the copy number level. Genomic copy number variations in single cells classified as epithelial in both the initial (P0) sample and in established tumoroids were estimated using the inferCNV package^18^, with immune cells from the original samples used as the reference cells during analysis (**Figure 3i, Supplementary Fig. S5**). Epithelial cells in the P0 samples had distinct regions of copy number gain or loss, which were largely conserved in tumoroid cells (**Figure 3i, Supplementary Fig. S5**). Gene expression levels for the subset of genes used in the inferCNV analysis were highly correlated between the initial epithelial subpopulation and the established tumoroids, with Pearson’s correlation coefficient values of *r*=0.77 and *r*=0.75 for HuCo3209 and HuCo021320, respectively (**Supplementary Fig. S5**). In support of these findings, copy number estimates for genes covered by targeted DNA sequencing showed high levels of correlation between initial tumors and tumoroid cultures at the bulk sequencing level (**Supplementary Fig. S5**).

### Long-term expansion of tumoroids in suspension culture

To evaluate the utility of patient-derived tumoroid cultures as long-term in vitro models, cryopreserved cultures derived from multiple colorectal and lung donors were thawed and maintained for up to 50 passages, with the cultures that were first established undergoing the longest evaluation. Experiments focused on suspension culture in non-tissue culture treated flasks with a low concentration of BME proteins (2% v/v) to provide ECM cues, a technique that facilitated routine maintenance of tumoroid lines. Patient-derived tumoroid cultures maintained their donor-specific morphologies during long-term suspension culture (**Figure 4a**). Tumoroid density in these representative images should not be taken as an indicator of growth, as the cells floating in the suspension culture can drift. Instead, cell growth rate was calculated by dissociating tumoroids to single cells during passaging and reseeding a known number of cells. Cell doubling time was donor-dependent and averaged around 65 hours for colorectal tumoroids – on par with that of 2D colorectal cancer cell lines – and 90-100 hours for lung tumoroids. The established tumoroids grew steadily for over 40 population doublings (**Figure 4b**). Testing included an additional cryopreservation and recovery step (indicated by arrows) and was concluded based on timing limitations, with no indication that cells would not expand further. Additionally, proof-of-concept experiments indicated that tumoroid lines expanded in suspension could be tumorigenic, with two of the lines tested (HuCo1044 and HuLu051921) forming tumors in NSG mice 40-80 days after subcutaneous injection of 3e6 cells/mouse. In both cases, resulting tumors showed histological similarity to the tumoroid cultures (**Supplementary Fig. S6**).

**Figure 4.**
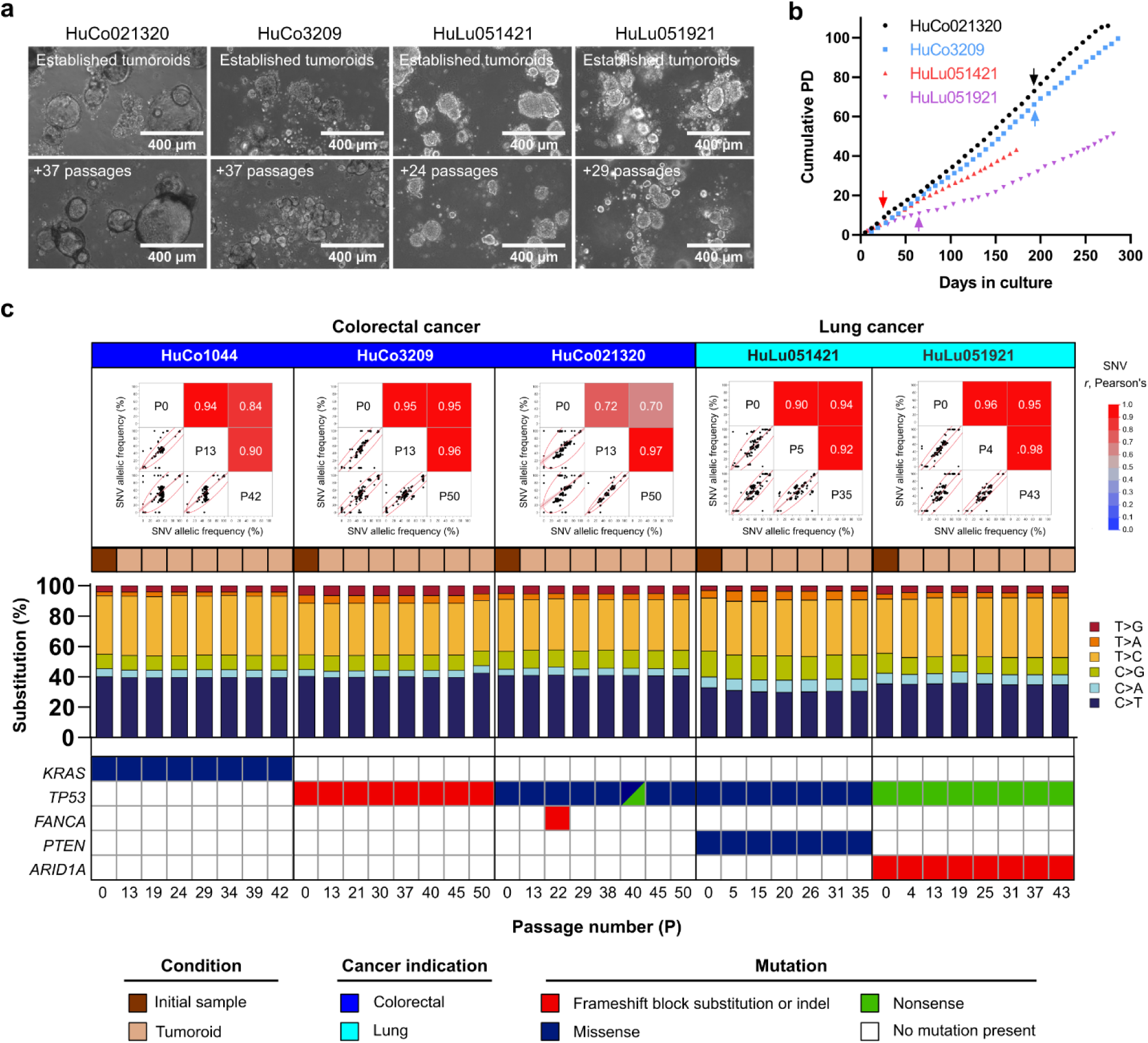
OncoPro medium-derived tumoroids are stable during long-term culture. (**a**) Morphology and (**b**) growth rate of tumoroid lines during long-term culture. Cumulative population doublings (PD) were measured by cell counts at each passage. Tumoroid lines were recovered from cryopreservation at the beginning of this experiment (day 0), and additional cryopreservation and recovery points are indicated by arrows for each culture. (**c**) Pearson’s correlation of single nucleotide variants (SNVs) between tumor samples (P0) and tumoroids at bank and at late passage numbers (**top**). Each dot represents the variant allelic frequency (VAF) for 1 genetic locus covered by the Oncomine™ Comprehensive Assay v3 (OCAv3). Single base substitutions (**bar graphs, middle**) and oncogenic mutations (**heat maps, bottom**) from multiple time points during this study. The oncogenic mutations were called using the Oncomine™ Variants filter 5.18 within the Ion Reporter™ Software.

Genomic stability was assessed every 5-10 passages during suspension culture by characterizing patient-specific single nucleotide variants (SNVs) using targeted sequencing via the OCAv3 assay. For each culture, the allelic frequency of identified SNVs were compared between uncultured tumor cells (P0) and early- and late-passage cultures (**Figure 4c, top**). Linear fits of this data show that SNV frequency is highly correlated (Pearson’s correlation coefficient) during long-term culture. In particular, high correlation coefficients (*r* ≥ 0.90) were observed between early and late passage cells, suggesting minimal drift over time in culture after tumoroid line establishment (**Figure 4c, top**). To understand the generalizability of this finding, a tumoroid line (HuCo021320) was transferred to another site and expanded from cryopreservation for >20 passages by different users. Similar correlation values were observed in SNVs (**Supplementary Fig. S7**). Further analysis of single base substitutions showed conserved patterns of substitution percentages and transition-transversion mutations between tumor tissue and matched tumoroid cultures during extended *in vitro* propagation (**Figure 4c, middle**). Finally, known driver mutations were well conserved from tumor tissue across multiple passages in tumoroid culture (**Figure 4c, bottom**), with uncommon instances of new mutations being called, some of which were not called again in sequencing runs of later passage cultures.

Transcriptome stability in patient-derived tumoroid cultures was first assessed using bulk RNA sequencing. Samples from multiple passages clustered by donor during principal component analysis (PCA; **Figure 5a**), and gene expression levels clustered by donor in unsupervised hierarchical clustering analysis of Euclidean distances between the samples (**Supplementary Fig. S8**). Similarly, tumoroid growth rates and gene expression levels were conserved for a given donor across different medium production lots (**Supplementary Fig. S7**). Differential gene expression analysis of 2 colorectal tumoroid donors at early (P13) vs. late (P50) timepoints demonstrated no significant change in >97% of genes (**Figure 5b**). Pathway analysis of differentially expressed genes (DEGs) in either early- or late-passage tumoroids revealed two significantly enriched KEGG pathways relating to protein translation and metabolism in late-passage tumoroids, with no KEGG pathways relatively enriched in early-passage tumoroids (**Figure 5b**). Similarly, comparison of early passage (P4 or P5) to late passage (P37 or P35, respectively) lung tumoroids revealed that only 1.1% of genes were differentially expressed, with enrichment for a small number of KEGG pathways primarily related to inflammatory and neural signaling in early-passage tumoroids and no significantly upregulated pathways at later passages (**Figure 5b**). Having demonstrated the maintenance of overall gene expression patterns during long-term culture, we next asked if the clinical subtype of tumoroids was maintained. We used a gene expression-based PAM38 classifier^19^ to classify colorectal cancer tumoroids into 5 consensus molecular subtypes: enterocyte, goblet-like, inflammatory, stem-like, and transit amplifying. Tumoroids from different donors represented unique subtypes (**Figure 5c**), indicating that diverse phenotypes can be maintained in tumoroid culture. HuCo021320 tumoroids resembled goblet and inflammatory subtypes; HuCo03209 and HuCo031721 tumoroids most closely matched the transit amplifying subtype, with a small proportion of stem-like cells; HuCo111622 cells were mostly enterocyte or goblet-like; and HuCo1044 tumoroids contained mixed subtypes (**Figure 5c**). Importantly, all the donors used in this analysis represented their respective clinical subtypes when cultured for 10-37 passages. Characterization of HuCo021320 colorectal cancer tumoroids by scRNA-seq showed that passage 27 tumoroids overlapped with passage 10 tumoroid samples and were distinct from those from another donor (**Figure 5d**), demonstrating maintenance of identity at the single-cell level. Additionally, CNV estimates inferred from scRNA-seq data (**Figure 5e**) or estimated from targeted genomic sequencing (**Supplementary Fig. S9**) indicated that tumor-specific ploidy values were maintained from early- to late-passage tumoroids.

**Figure 5.**
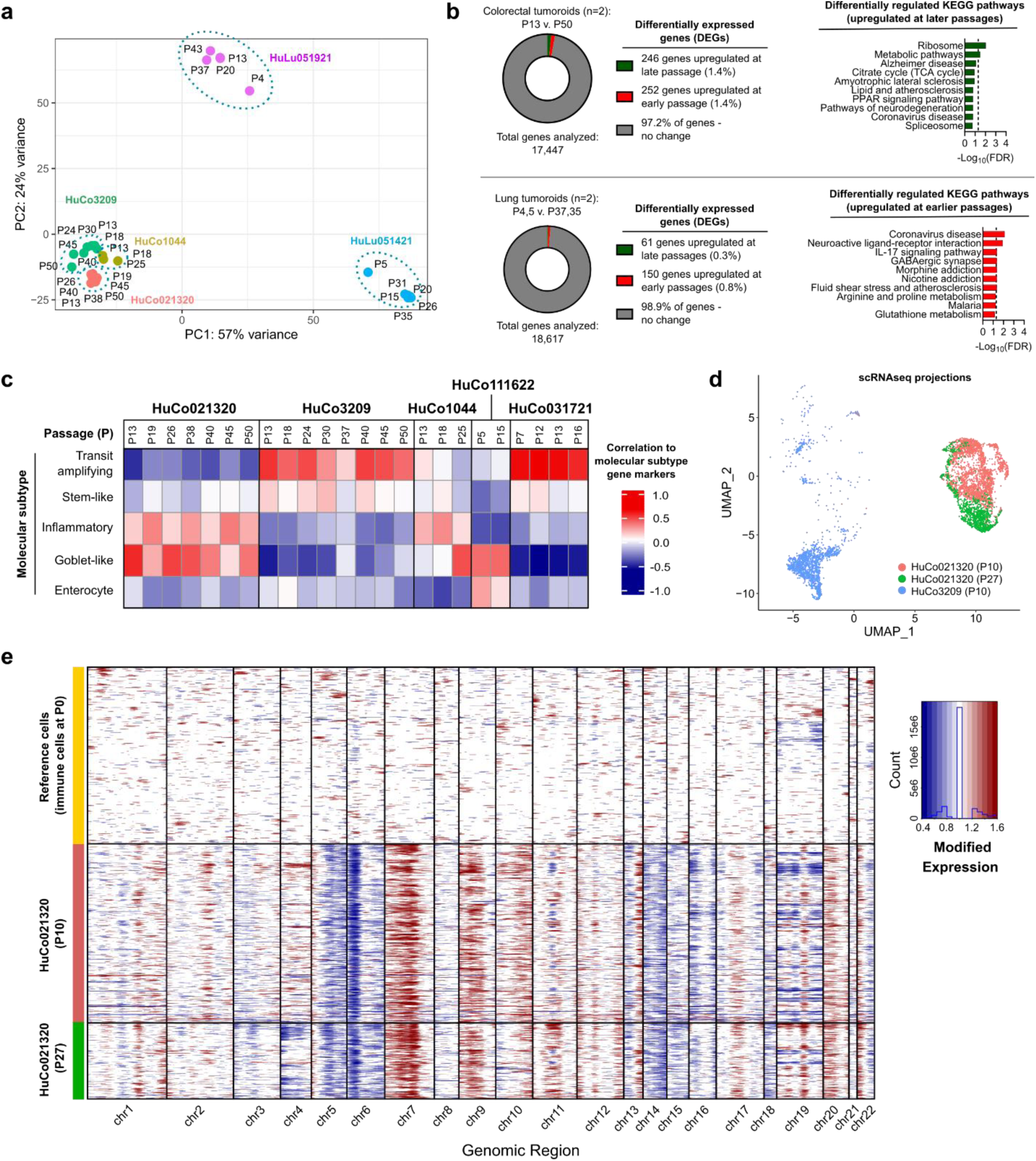
OncoPro medium-derived tumoroid transcriptomes are stable during long-term culture. (**a**) Principal component analysis (PCA) of global gene expression patterns in human colorectal (HuCo) and lung (HuLu) tumoroid cultures over multiple passages (P). (**b**) Differential gene expression (DEG) analysis of early-versus late-passage tumoroids, with DEGs called at fold change>2 and false discovery rate (FDR)<0.05, for HuCo and HuLu samples. Pathway analysis ofDEGs revealed few significantly enriched Kyoto Encyclopedia of Genes and Genomes (KEGG) pathways (FDR<0.05) in late-passage colorectal tumoroids and early-passage lung tumoroids; the top 10 KEGG pathways for colorectal and lung tumoroids are shown. No KEGG pathways were enriched in early-passage colorectal tumoroids or late-passage lung tumoroids. Dotted line indicates FDR=0.05. (**c**) Consensus molecular subtypes from gene expression analysis of colorectal patient-derived tumoroids through multiple passages. (**d**) Uniform manifold approximation and projection (UMAP) for dimension reduction plot of 4,905 total cells for HuCo021320 (early, P10; and late, P27) and HuCo3209 (early, P10) tumoroid cultures. HuCo3209 was used as a control donor for comparison. (**e**) Single cell copy number variation (CNV) plots for HuCo021320 (n=4,973 genes, by column) of early (P10) and late (P27) passage tumoroid cultures. Each row represents one cell. InferCNV was used to estimate CNV in epithelial cells, using immune cells present in dissociated tumor tissue (P0) as reference.

### Growth and genotypic stability of previously established tumoroid cultures in OncoPro medium

To test the generalizability of the culture system with publicly available tumoroid models, we next tested the expansion of tumoroid cells from the United States National Cancer Institute Patient-Derived Models Repository (NCI PDMR)^20^, and an additional, commercially available colorectal tumoroid line (3dGRO™ ISO72, Millipore Sigma). These models come with recommendations for use of culture media that is generated by combining basal medium with various supplements, growth factors, and, at times, conditioned media, which we collectively refer to as “homebrew” tumoroid media. Models tested were procured as cryopreserved cells. In our experiments, cells were plated into BME-embedded culture in source-recommended media (NCI PDMR recommended homebrew media or vendor-recommended homebrew medium; see Materials and Methods and **Supplementary Table S5** for details and modification of NCI PDMR media recipes to incorporate recombinant proteins), or in OncoPro medium in either embedded or suspension formats (with indication-specific growth factors and supplements as required; see **Supplementary Table S1**) in parallel (**Figure 6A**). Testing was performed either upon receipt of cryopreserved material or after expansion of a tumoroid bank from initial NCI PDMR material using modified NCI PDMR media recipes (see details in **Supplementary Table S5 and Supplementary Table S6**). Growth of cells was monitored over time by obtaining cell counts at each passage, reseeding a known number of cells, and calculating cumulative population doublings. Additionally, cells were sampled for DNA and RNA purification from each condition upon culture initiation and after 3-6 passages post-thaw for most cultures (see **Supplementary Table S6**). In some instances, low initial cell yield precluded the sampling of cells at the time of initial thaw for RNA sequencing. Testing was performed using 4 colorectal tumoroid lines, 4 lung tumoroid lines, 4 breast tumoroid lines, 3 head and neck tumoroid lines, and 3 pancreas tumoroid lines (**Supplementary Table S6**). No crypt villus organoids or colon crypt organoids were observed as only carcinoma cells were procured for this study.

**Figure 6.**
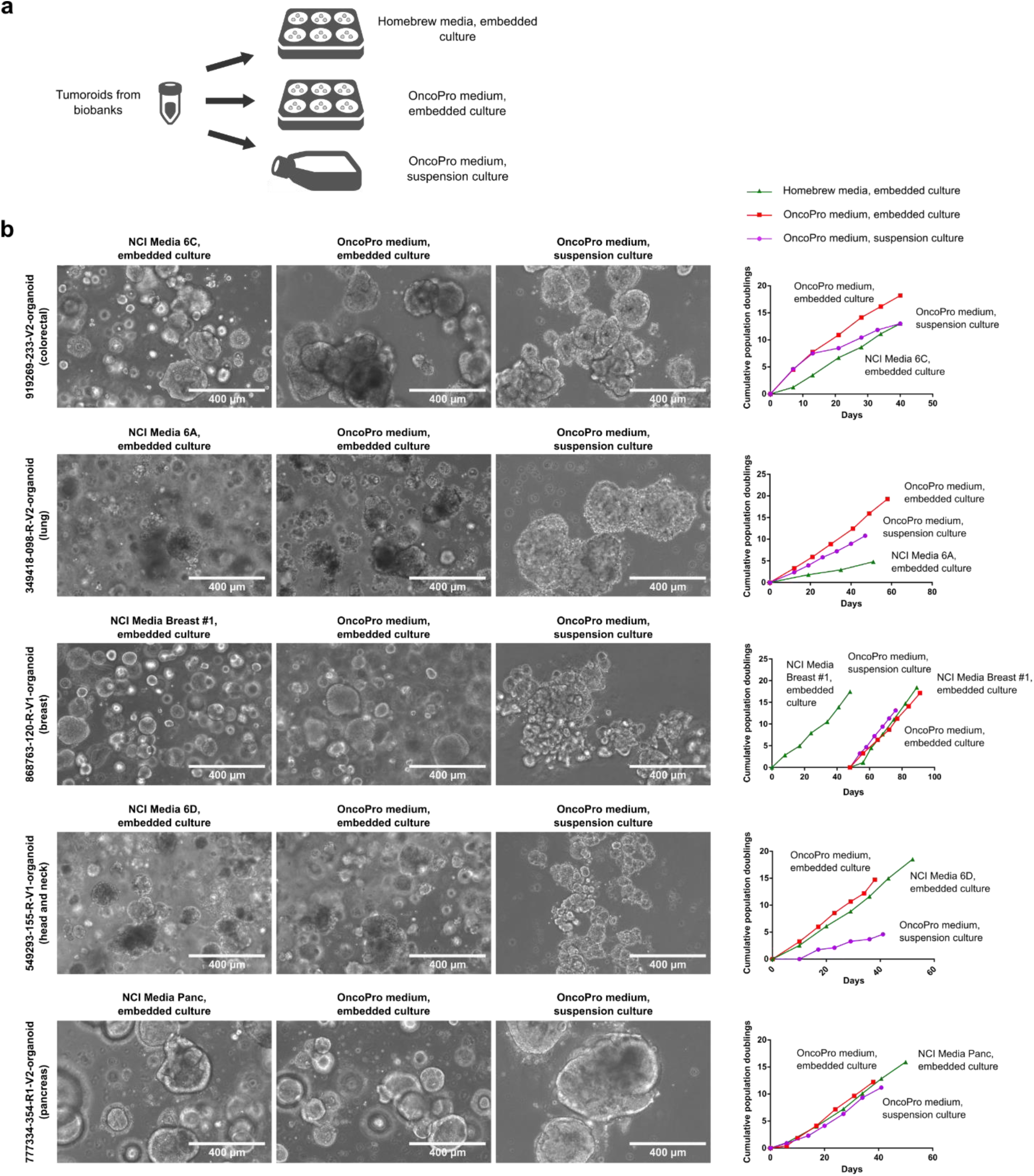
Publicly available tumoroid lines can adapt well to culture in OncoPro medium. (**a**) Outline of study comparing growth rate of tumoroids from biobanks (for example, National Cancer Institute Patient-Derived Models Repository, PDMR) in homebrew medium to growth of tumoroids in OncoPro medium in embedded or suspension culture formats. (**b**) Tumoroid lines from colorectal, lung, breast, head and neck, and pancreas cancer cultured in PDMR-recommended homebrew media or in OncoPro medium in embedded or suspension culture have similar morphologies and growth rates. For model 868763-120-R-V1-organoid, cells were initially expanded in PDMR-recommended media for several passages to create a working bank and then cultured in all three conditions in parallel. Scale bar = 400 µm.

In our testing, morphologies and growth rates of tumoroid cultures were similar across media systems and culture methods (**Figure 6b,c; Supplementary Fig. S10**). In some cases, tumoroid growth rate was higher in embedded culture than in suspension culture, and vice versa. Growth rates were generally similar between the NCI PDMR or vendor recommended medium and OncoPro medium, with some models exhibiting a slight growth advantage in one system over the other. For some tumoroid lines, cells from one arm of the study were lost during culture or were not fully tested in the suspension culture format (denoted as “Data not available”; see also **Supplementary Table S6**). One NCI PDMR lung cancer line, LG0481, did not expand in OncoPro medium but grew in NCI PDMR Media 6A, while a head and neck cancer tumoroid line, 832693-133-R-V1-organoid, expanded in OncoPro medium in embedded and suspension formats but not in the recommended NCI PDMR Media 6D (**Supplementary Fig. S10**). To test whether a change in media system or culture format was associated with genetic or transcriptional drift in these models, we performed targeted mutational profiling and transcriptional analysis to compare initial cells (as received, or cells from an internally established cell bank using NCI PDMR media; see **Supplementary Table S6** for details) to cells expanded in various arms of the study. Tumoroid cultures from different donors and cancer indications harbored a heterogeneous set of cancer driver genes affected by missense mutations, non-sense mutations, or frameshift mutations (**Figure 7a**). In general, oncogenic mutations were preserved between culture conditions, though some mutations were detected in initial material that were not called in one or more expansion arms, and vice versa (**Figure 7a**). No clear pattern of loss or emergence of mutations following culture in a given media system or culture format was noted, and SNV presence and variant allelic frequency were highly concordant for a given tumoroid line (*r*>0.97 when all samples were batched and compared) regardless of the system in which they were grown (**Figure 7a, Supplementary Table S7**). The minimal cross-culture correlations observed in this data indicate retention of donor specificity (**Supplementary Table S7**). SNV concordance between initial cells and tumoroids expanded in each arm of the study was also high (>97% for 47/48 of comparisons), indicating that culture conditions were not driving the loss or emergence of SNVs (**Supplementary Fig. S11**).

**Figure 7.**
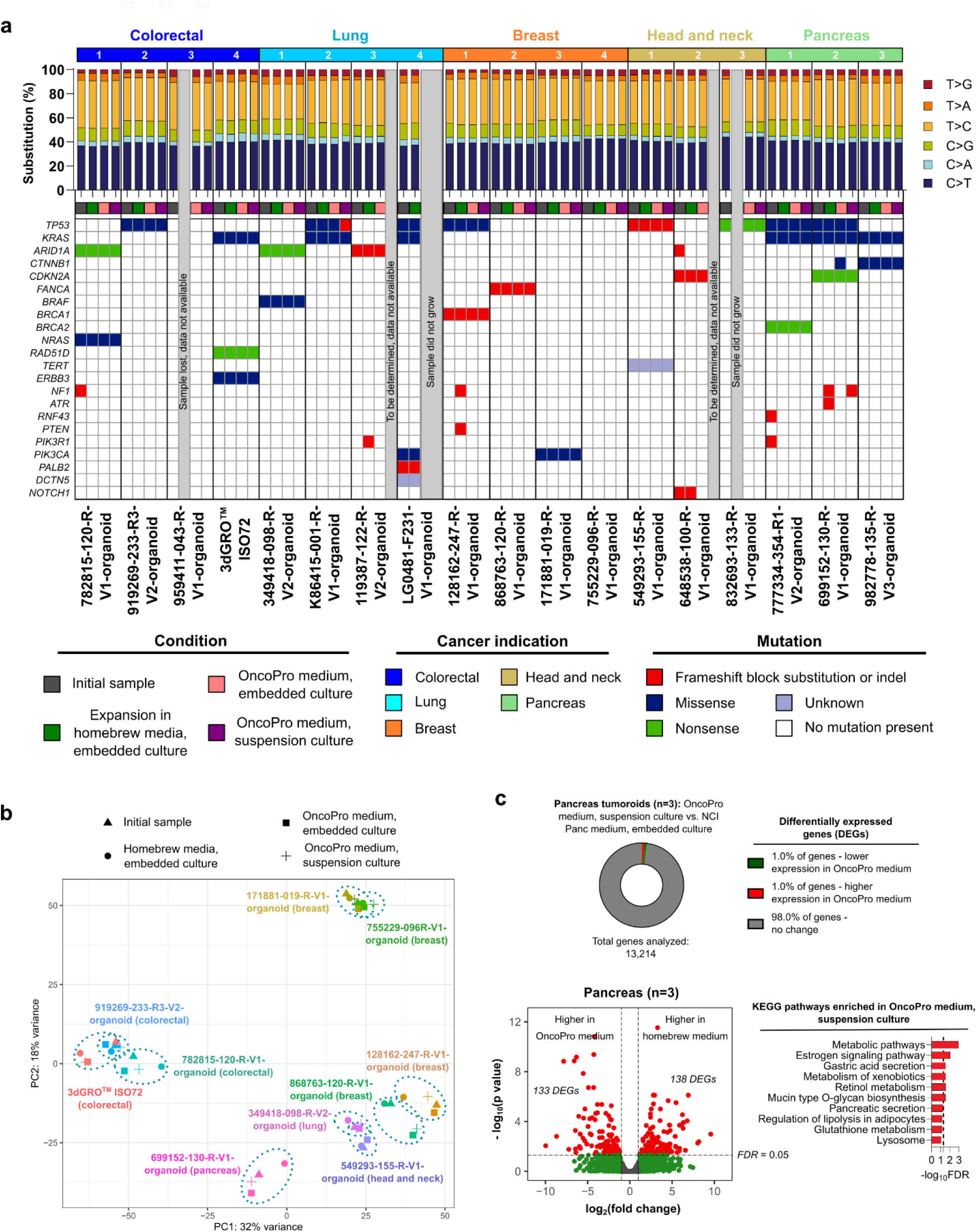
Previously established tumoroid lines adapted to OncoPro medium have similar genomic and transcriptomic characteristics compared to the cultures grown in homebrew media. (**a**) Single nucleotide base substitutions (**top**) and oncogenic single nucleotide variants present in the samples following sequencing with the Oncomine™ Comprehensive Assay v3 (OCAv3) and mutation calls called using the Oncomine™ Variants filter 5.18 within the Ion Reporter™ software (**bottom**). (**b**) Principal component analysis (PCA) plot comparing gene expression levels in initial cells (triangles; directly from vendors or from internal cell banks) and cells expanded using homebrew embedded culture (circles), OncoPro medium embedded culture (squares), and OncoPro medium suspension culture (crosses). (**c**) Differential gene expression analysis of pancreas tumoroids cultured in PDMR-recommended homebrew media (embedded format) and OncoPro medium (suspension format; n=3), showing differentially expressed genes (DEGs) at fold change>2 and false discovery rate (FDR)<0.05. Percentage change in transcriptome (**c**, **top**) and analysis for enriched Kyoto Encyclopedia of Genes and Genomes (KEGG) pathways in genes highly expressed during suspension culture in OncoPro medium (**c**, **bottom**). Top 10 results are shown. Dotted line indicates FDR=0.05. No KEGG pathway was enriched when highly expressed genes in homebrew medium were used in the analysis.

PCA of gene expression profiles from initial samples (received from vendor or from initial cell bank) versus those expanded in this study showed clustering by donor rather than by growth condition, indicating maintenance of gene expression profiles in OncoPro medium (**Figure 7B**). In some instances, low initial cell yield precluded the sampling of cells at the time of initial thaw for RNA sequencing; including samples in which we could not obtain an initial gene expression profile led to similar clustering by PCA after bulk RNA sequencing (**Supplementary Fig. S12**). To understand what differences are driven by the suspension format, we compared 15 pairs of suspension and embedded tumoroid cultures (n=3 colorectal, n=4 lung, n=2 pancreas, n=2 head and neck, n=4 breast) maintained in OncoPro medium at similar passage numbers. In this comparison, only 1% of genes were differentially expressed (**Supplementary Fig. S12**). Genes upregulated in embedded culture were related to stress and hypoxic response, possibly due to the relatively higher cell-cell proximity within these cultures compared to the suspension format.

On an indication-specific basis, tumoroids expanded in OncoPro medium suspension culture and in homebrew media embedded culture were further compared to elucidate differences driven by the suspension culture approach. For pancreas cancer samples, only 2% of genes were differentially expressed between these conditions (**Figure 7C**). KEGG analysis showed that genes highly expressed during suspension culture in OncoPro medium (compared to embedded culture in NCI Panc medium) were associated primarily with metabolic signaling pathways (**Figure 7C**). No KEGG signaling pathways were enriched when analyzing genes upregulated in NCI Panc medium. Similar results were obtained for colorectal, lung, and breast cancer tumoroids, where expression levels for <4% of genes were significantly different when comparing tumoroids expanded in homebrew embedded culture to those grown in OncoPro medium in the suspension format (**Supplementary Fig. S13**).

Clinical subtypes based on gene expression levels were also generally retained across media systems. Classification of colorectal cancer tumoroids using the PAM38 gene classifier showed that original and OncoPro medium cultures of the 782815-120-R-V1-organoid, 919269-233-R3-V2-organoid, and 3dGRO™ ISO72 tumoroids maintained the transit amplifying, goblet-like/inflammatory, and transit amplifying/stem-like subtypes present in the original samples, respectively, after culture in OncoPro medium (**Supplementary Fig. S13**). Similarly, the basal subtype of triple negative 128162-247-R-V1-organoid and 868763-120-R-V1 breast cancer tumoroids (by the PAM50 gene signature) was consistent across culture formats (**Supplementary Fig. S13**). However, PAM50 subtyping of 171881-019-R-V1-organoid and 755229-096-R-V1 lines showed that OncoPro cultures adopted HER2 subtypes, while the original cultures had characteristics of luminal B cancers. These lines are positive for estrogen receptor (ER) and progesterone receptor (PR) expression (ER+/PR+), and we noted that the expression of estrogen receptor 1 (ESR1) and progesterone receptor (PGR) genes was lower in these cultures after transition to OncoPro medium. Of note, the expression of these genes was still significantly higher in the OncoPro medium-transitioned ER^+^/PR^+^ lines when compared to the triple-negative lines, indicating retention of ER^+^/PR^+^ expression in OncoPro medium (**Supplementary Fig. S13**). Other groups have shown some loss of these receptors in 30% of conditions^21^, and further optimization of medium conditions may be required for maintenance of ER/PR levels that more consistently reflect those observed in clinical samples. Overall, these data indicate that OncoPro medium can be used for the culture of existing tumoroid models from colorectal, lung, breast, head and neck, and pancreas cancers, with minimal expected changes in growth rate, mutational profile, or gene expression levels.

### Tumoroid suspension cultures are amenable to medium-throughput, multiplexed immune cell-mediated cytotoxicity assays

To demonstrate the compatibility of tumoroids with genome engineering and cytotoxicity assays, we engineered a colorectal tumoroid line, HuCo1044, to express green fluorescent protein (GFP). After dissociation and exposure to lentivirus in suspension culture, 67% of tumoroid cells were GFP-positive (GFP+), which increased to over 99% after antibiotic selection (**Supplementary Fig. S14, Supplementary Video S1**). Targeted genomic sequencing and transcriptome profiling indicated that the engineered cells had a high degree of similarity to the parental culture (**Supplementary Fig. S14**). To simulate immune-oncology development workflows, we co-cultured GFP+ tumoroids with primary natural killer (NK) cells or an immortalized NK cell line, NK-92. Primary NK cells isolated from peripheral blood mononuclear cells via negative selection demonstrated 94.8% purity (CD45+ CD56+), were not enriched for CD3+ T cells (0.25%), and contained 33.3% CD16^bright^ cells (**Supplementary Fig. S14**). Tumoroids (target cells) were passaged and seeded alone, and their expansion could be tracked by GFP signal (**Figure 8a,b)**. After 64 hours, NK cells (effector cells) were seeded at various effector to target (E:T) ratios. Co-incubation of tumoroids and NK cells led to a dose-dependent decrease in tumoroid GFP signal and a concomitant increase in caspase 3/7 activity (**Figure 8a,b**). Quantitative image analysis demonstrated that tumoroid-only control cells continued to grow, leading to an increase in GFP signal in the absence of caspase activity, while the presence of NK cells led to decreasing GFP intensity in a dose-dependent manner (**Figure 8b**). Similar results were demonstrated with the NK-92 cell line, though its killing efficiency appeared lower than that of the primary cells (**Figure 8b**). In summary, tumoroids cultured in OncoPro medium were compatible with genetic engineering and could be utilized to demonstrate the effectiveness of cancer-targeting regimens.

**Figure 8.**
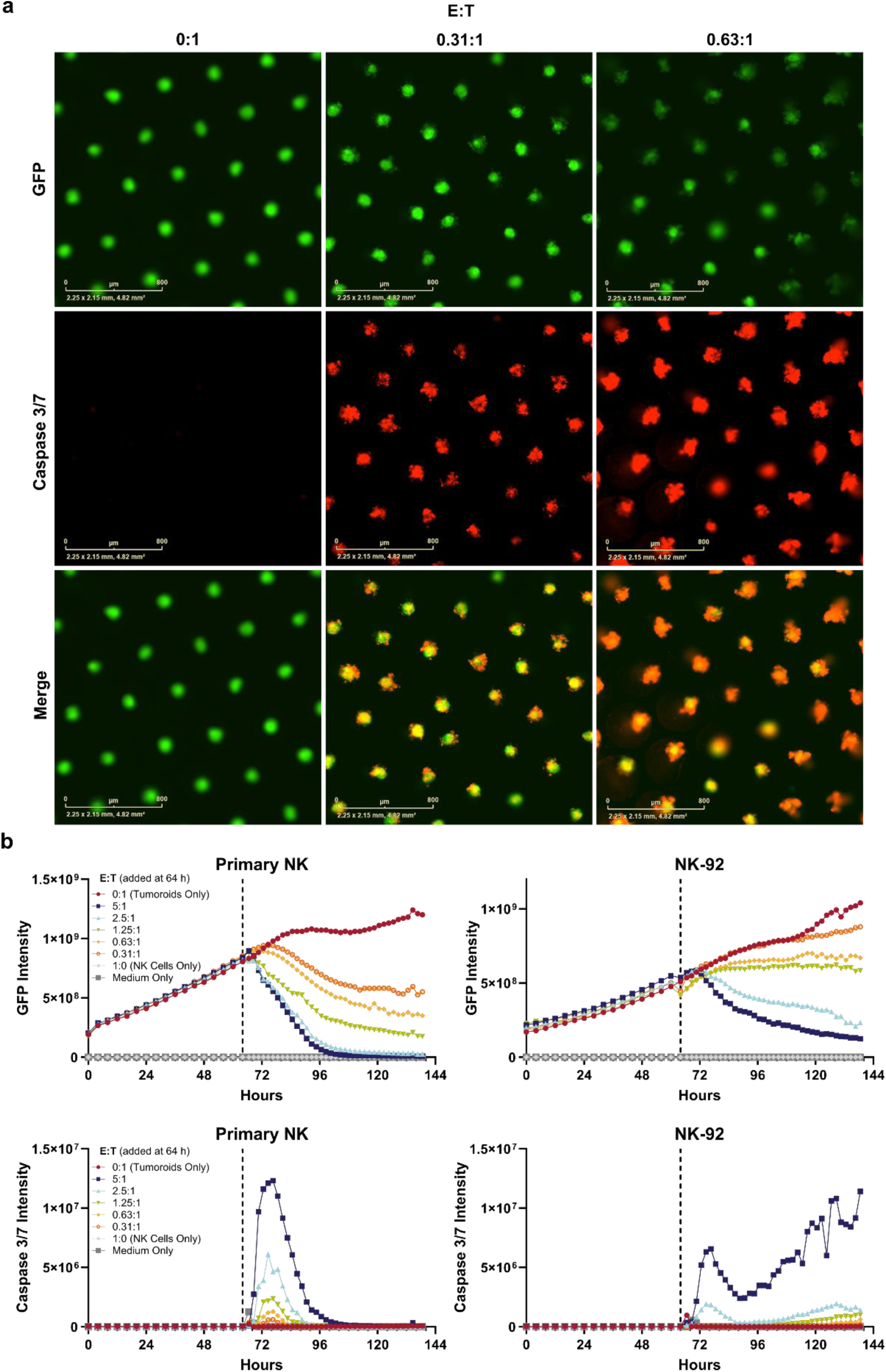
Tumoroid suspension cultures are amenable to multiplexed assays for immune cell cytotoxicity. A colorectal tumoroid line stably expressing green fluorescent protein (GFP) was dissociated and seeded into 96 well microcavity plates. After 64 hours, natural killer (NK) cells from a primary source or the NK-92 cell line were added at various effector to target (E:T) ratios along with an indicator of caspase 3/7 activity, Invitrogen™ CellEvent™ Caspase-3/7 Red. (**a**) Representative images of HuCo1044-GFP cells acquired on an Incucyte^®^ Live-Cell Analysis System 24 hours after addition of primary NK cells at multiple E:T ratios (0:1 represents tumoroids only condition). Scale bar = 800 µm. (**b**) Quantitative image analysis of green (HuCo1044-GFP) or red (caspase 3/7 activity) integrated fluorescence intensity (green or red calibrated units × µm^2^/well) before and after addition of Primary NK or NK-92 cells at 64 hours (dotted line).

## Discussion

Here, we demonstrate the utility of a novel tumoroid culture medium, Gibco OncoPro Tumoroid Culture Medium, for the culture of patient-derived 3D tumoroid models. The system was used for both (1) deriving novel tumoroid lines from cryopreserved dissociated tumor cells and fresh surgical resection samples, and (2) culturing established cancer organoid lines. When supplemented with tissue-specific niche factors, we demonstrated the utility of OncoPro medium for culturing tumoroids from a variety of solid tumors, including colorectal, lung, pancreatic, head and neck, endometrial, and breast cancers. During culture, mutations and gene expression signatures characteristic of the starting material were retained (**Figures 2, 3, 7**), and select lines tested were tumorigenic (**Supplementary Fig. S6**). Furthermore, tumoroids from a wide range of tumor stages, grades, mutational backgrounds, molecular subtypes, and donor demographics were supported (**Figure 2, 7; Supplementary Table S2**). There is not a clear pattern of these variables that determines compatibility with the medium, though this bears monitoring as sample size increases. We expect that supporting niche factors that enable culture from additional cancer indications will be uncovered, and additional efforts using both fresh tissue resections and existing tumoroid models from the NCI PDMR are in progress.

Tumoroid culture establishment represents a key technology for enabling patient-derived cancer models and is more affordable than establishing novel patient-derived xenograft (PDX) models^22^. In our studies, the success rate for stable tumoroid line establishment varied depending on multiple factors, but, with an optimized protocol, remained around 50% for colorectal and endometrial cancers. Our initial studies suggest that this success rate is lower in lung and breast cancers. We noted much higher success rate (>85%) of tumoroid formation during short-term culture across indications, as also observed by others^23,24^. Additional protocol improvements or tissue-niche factors may further enhance derivation rates; others have demonstrated that derivation using multiple bespoke homebrew media in parallel can further enhance success^25^.

Altogether, tumoroids retain the major genomic mutations of the tumors from which they were derived. Targeted genomic profiling to compare tumor material to matched tumoroid cells revealed occurrence of oncogenic mutations and donor-specific SNVs at similar allelic frequencies in both sets. Copy number variations were also maintained, as estimated broadly over the genome using scRNA-seq and inferCNV or by the OCAv3 assay. Mutations identified in tumor and tumoroid samples will depend on the sequencing method and analysis pipeline, and broader genomic sequencing would supplement the targeted approach utilized here. Comparison of epigenetic modifications to tumor cells would also be worthwhile given recent findings that tumoroids retain greater sensitivity to epigenetic vulnerabilities than 2D cell lines^26^. The loss of expression of immune and other stromal cell signaling pathways during derivation in OncoPro medium represented the major changes in the gene expression profiles of tumoroid cultures when compared to primary tumor material. Other media systems have also demonstrated loss of signaling associated with non-malignant cell components over the course of weeks in culture^27^. At the single-cell level, comparison of malignant tumor cells and tumoroid cells demonstrated that transcriptomes were similar, with a correlation coefficient of 0.85, further indicating that loss of non-tumor cell populations drive much of the observed transcriptional changes observed during culture establishment. Transcriptional changes within the tumor cell population may be caused by removal of cell-cell contacts or paracrine signaling from non-malignant cells. Preservation or re-addition of other cell types in co-culture or using *in vivo* models may restore these cues^28,29^.

As compared to several other published methods^11,13,21,30–32^, our proprietary and novel serum-free medium does not require conditioned medium or Wnt agonists such as R-spondins. The lack of Wnt activation may protect against non-malignant cells outcompeting malignant cells in tumoroid line derivation, a known issue with some cancer indications, including lung and breast cancers^14,16,33^. Despite lacking Wnt agonists, OncoPro medium supported multiple tumoroid models from the NCI PDMR that are typically grown in systems containing Wnt or R-Spondins. This is in line with literature suggesting that exogenous Wnt activation may not be required for in vitro growth of multiple cancer indications^25,34,35^, either by gene mutation, endogenous production, or through transcriptional programming. In line with this, we verified the presence of APC mutations^17^ in select colorectal tumoroids (**Supplementary Table S4**). In addition to Wnt agonists, other factors and small molecules demonstrated to be necessary for the growth of non-malignant organoids are present in many published tumoroid media formulations^21,32,34^.

Once established, comparison of early and late passage tumoroid cultures indicated stable growth rates, with retention of CNV levels and SNV presence and frequency, demonstrating stability over months in culture. This result held for both internally-derived tumoroid lines and tumoroids procured from external parties, where OncoPro medium supported commercially available tumoroid models (17/18 tested here). Testing revealed the importance of beginning with sufficient cell number from high-quality cryobanks or ongoing cultures, and passage of individual cultures when tumoroids reached sufficient size to prevent overgrowth (even if it meant passaging different media conditions at different times). Overall, growth and morphology were similar between homebrew media systems and OncoPro medium embedded or suspension cultures, with occasional donor-to-donor variability in growth rate between culture formats or media. One lung adenocarcinoma line (LG0481-F231-V1-organoid) expanded in the NCI recommended medium but not in OncoPro medium, while one lip/oral cavity squamous cell carcinoma (head and neck) sample (832693-133-R-V1-organoid) expanded in OncoPro medium but not the NCI recommended medium in our hands. NCI PDMR standard protocols rely on the addition of fetal bovine serum (FBS) through addition of L-WRN conditioned medium, and it is possible that the instances in which tumoroid growth was slower in NCI PDMR media compared to OncoPro medium was influenced by our approach using recombinant Wnt (W), RSPO (R), and noggin (N) sources, which changed the source of WRN in the media and eliminated FBS from these cultures.

Critically, genomic mutations and gene expression patterns were maintained during culture of established tumoroid lines in OncoPro medium in both embedded and suspension formats. These comparisons also revealed that few genes were differentially expressed after months in culture. Molecular subtypes of colorectal and triple-negative breast cancers were maintained during tumoroid culture, suggesting that key clinical aspects of gene expression signatures are stable during culture using OncoPro medium. Such maintenance of clinical subtype during tumoroid establishment has been named as critically important to demonstrate the physiological relevance of in vitro cancer models, especially with regard to drug response^21,23,36^, and previous publications have mixed claims regarding the ability of tumoroids to adequately maintain subtypes representative of the tissue from which the tumoroid was derived or generate biobanks with tumoroid lines representing certain clinical subtypes. In line with this result, we observed some shift of hormone-receptor positive lines towards a HER2 subtype in OncoPro medium, with concomitant decreases in estrogen receptor and progesterone receptor gene expression levels compared to cryopreserved tumoroids received from the NCI PDMR (**Supplementary Fig. S13**). Molecular subtyping approaches are dependent on the timepoint for comparison (tissue versus early passage for later comparison), cancer type of interest, assay of choice (scRNA-seq versus bulk RNA sequencing), and analysis pipeline, to name a few, and can thus be difficult to compare across studies. Future testing of the functional impact of maintenance of molecular subtype by screening therapeutic compounds for early versus late passage organoids and, ultimately, a retrospective analysis of organoid response in vitro compared to primary tumor response would be more definitive.

We anticipate that standardized and easy-to-use tumoroid culture systems will promote their use in assays in which immortalized 2D cancer cell lines are commonly used. To this end, OncoPro medium was designed to enable consistent tumoroid media preparation and compatibility with multiple cancer indications and formats during 3D cell culture. The serum- and conditioned-medium free formulation streamlined medium preparation and allowed for standardization between users and institutions and from lot-to-lot (**Supplementary Fig. S7**), a clear need in the cancer organoid field^15^. Suspension culture is emerging as a more accessible alternative to embedded tumoroid culture, which requires greater amounts of expensive matrix material, precise pipetting, and, in many cases, careful control of reagent temperature. In previous studies, tumoroids exhibited reduced or minimal growth in suspension without ECM proteins, though the addition of 2-5% volume/volume ECM potentiated growth^24,37–39^. These studies have demonstrated suspension culture of primary colorectal, metastatic colorectal, and esophageal tumoroids, typically for 2-4 weeks, with some examples of longer durations (20-25 weeks)^39^. Another approach is to encapsulate tumoroid or organoid cells in small droplets of ECM that float in suspension medium, which has been demonstrated in NSCLC and non-malignant intestinal organoids^40,41^. In our study, the addition of 2% BME generated a reliable suspension culture system suitable for long-term culture across a variety of cancer indications when used in conjunction with OncoPro medium. The time required for laboratory personnel to subculture tumoroids cultured in suspension decreased by approximately 2-fold compared to the embedded approach in our testing, facilitating maintenance of tumoroid models for downstream assays. Importantly, we demonstrated few changes at the transcriptome level between tumoroids in embedded or suspension culture. Potential downregulation in hypoxia-related signaling, cytokine-cytokine receptor interactions, and TNFα signaling in suspension compared to embedded culture formats (**Supplementary Fig. S12**) should be considered during assay design.

As one example of how tumoroid assays could be implemented, we engineered a tumoroid reporter line to express GFP and used that line to assess NK cell cytotoxicity in a 3D culture format (**Figure 8**). Testing the cytotoxic potential on 2D models of solid tumor cancers does not adequately represent the physiological conditions with which NK cells will contend in vivo, namely dense tumor tissue and nutrient gradients^42,43^, which are better represented with tumoroid morphologies. However, NK cells are incompatible with embedded culture methods for tumoroids, as the BME inhibits NK cell penetration^44^, an issue which the suspension culture outlined here circumvents. Multiplexed reporters enabled analysis of NK-cell mediated cytotoxicity by two signals, GFP reduction and increased caspase-3/7 activity. These techniques could be readily adapted to research on other modalities of cancer therapeutics.

Finally, multiple reports have demonstrated that drug response in patient-derived tumoroid models, at least in some cases, correlates highly with matched donor response during clinical treatment^25,45–55^. Advancements in functional precision oncology - where measurable improvements in patient outcome are increasingly being cited - have been hampered by difficulties in standardized approaches for tumoroid culture^9,56–58^. Given the high degree of success of short-term culture in OncoPro medium and high fidelity of tumoroids to patient material demonstrated here, its use as a standardized cell culture medium in functional testing applications warrants further exploration. The high degree of site-to-site reproducibility observed in our experiments suggests that standardized cell culture approaches will improve the comparability of results from assays using patient-derived cancer models in the near term, helping to drive its adoption in both drug discovery and functional precision medicine applications.

## MATERIALS AND METHODS

### Tumor dissociation and initial cell seeding

Cryopreserved dissociated tumor cells or tumor resection samples were purchased from commercial vendors (Discovery Life Sciences, Huntsville, AL or Cureline Translational CRO, Brisbane, CA) following collection from patients of at least 18 years of age under informed consent and Institutional Review Board (IRB) approval. Tumor resection samples were shipped in Gibco™ Hibernate™-A Medium (Thermo Fisher Scientific, Waltham, MA, USA, cat# A1247-501) supplemented with, minimally, Gibco™ GlutaMAX™ Supplement (Thermo Fisher Scientific, cat# 35050-061), Gibco™ HEPES (Thermo Fisher Scientific, cat# 15630-206), Gibco™ penicillin-streptomycin (Thermo Fisher Scientific, cat# 15140-122), and Primocin^®^ (InvivoGen, San Diego, CA, USA, cat# ant-pm-05) at 4°C. Upon receipt, samples were immediately minced and washed. Samples were enzymatically dissociated for 1 hour at 37°C. Following dissociation, cells were counted using the Vi-CELL BLU Cell Viability Analyzer (Beckman Coulter, Indianapolis, IN, USA, cat# C19196) and plated at a concentration of 2.5×10^5^ viable cells/ml in Gibco™ OncoPro™ Tumoroid Culture Medium (Thermo Fisher Scientific, cat# A57012-01; supplemented as outlined in **Supplementary Table S1**) containing 10 µM Y27632 (Selleck Chemicals, Houston, TX, USA, cat# S1049) and 2% (v/v) Gibco™ Geltrex™ LDEV-Free Reduced Growth Factor Basement Membrane Matrix (Thermo Fisher Scientific, cat# A14132-02) in Nunc™ non-treated multidishes (Thermo Fisher Scientific). Additional dissociated cells were collected for subsequent RNA and DNA isolation. Samples for RNA analysis were collected by resuspending dissociated tumoroid pellets in Invitrogen ™ TRIzol™ Reagent (Thermo Fisher Scientific, cat# 15596-026) and storing at -20°C until RNA isolation. Dissociated cells for DNA profiling were pelleted, washed twice with DPBS with calcium and magnesium (DPBS (+/+); Thermo Fisher Scientific, cat# 14040-133), and pelleted again, with the cell pellet stored at -80°C until the time of DNA isolation. When sufficient tissue was available, minced tissue (pre-dissociation) was also collected for subsequent RNA isolation by submersion in TRIzol reagent, mechanical grinding with a pestle, and storage at -20°C, and for subsequent DNA isolation by flash freezing in liquid nitrogen and storing at -80°C. In general, P0 samples were cultured in suspension for 2-7 days prior to collecting cells and embedding in Geltrex matrix for several passages (details on embedded culture below; see also **Figure 1**).

### Tumoroid culture

Tumoroids were cultured in OncoPro medium as instructed by the supplier. Briefly, dissociated tumoroids, either at the time of thaw or following dissociation from tumor material, were resuspended in supplemented OncoPro medium containing 10 µM Y27632. For embedded culture in BME domes, cells were mixed with Geltrex matrix and OncoPro Tumoroid Culture Medium such that 50 µl of the final mixture contained 50,000 viable cells with a final Geltrex matrix concentration of 10 mg/ml. Geltrex matrix domes (50 µl/dome) were spotted on tissue culture plates before inverting plates and incubating for 30 min at 37°C and 5% CO_2_ to allow for Geltrex matrix polymerization. Plates were returned to the normal orientation and overlaid with supplemented OncoPro medium containing 10 µM Y27632. For suspension culture, dissociated cells were seeded at concentrations of 0.125-0.3×10^6^ cells/mL in supplemented OncoPro medium containing 10 µM Y27632 in non-tissue culture treated plates or flasks, after which 2% v/v Geltrex matrix was added and mixed. Medium was exchanged every 2-3 days for both culture formats following protocols in the OncoPro medium user guide.

Routine monitoring of tumoroid size was performed with an Invitrogen™ EVOS™ XL Core Imaging System (Thermo Fisher Scientific, cat# AMEX1000) or Invitrogen™ EVOS™ M7000 Imaging System (Thermo Fisher Scientific, cat# AMF7000). Tumoroids were subcultured when tumoroid diameter reached 100-300 µm, usually every 7-10 days (**Supplementary Figure S2**). Tumoroids were collected, and culture vessels were washed with cold Gibco™ DMEM/F-12, GlutaMAX™ supplement (Thermo Fisher Scientific, cat# 10565-018), which was added to the cell solution. Tumoroids were washed with cold DPBS without calcium or magnesium (DPBS (-/-); Thermo Fisher Scientific, cat# 14109-144), after which the cell pellet was resuspended in room temperature Gibco™ StemPro™ Accutase™ Cell Dissociation Reagent (Thermo Fisher Scientific, cat# A11105-01) supplemented with 10 µM Y27632. Tumoroids were incubated in Accutase reagent for 10-20 minutes at 37°C and 5% CO_2_ with periodic agitation. Tumoroids were then triturated 20 times with a P1000 pipet to mechanically disrupt the 3D structures. Dissociated tumoroids were counted and plated as described above. Dissociated cells were sampled for subsequent RNA and DNA isolation at various passages during culture. Additionally, cell banks were generated throughout culture by resuspending dissociated tumoroids at 2×10^6^ cells/ml in Gibco™ Recovery™ Cell Culture Freezing Medium (Thermo Fisher Scientific, cat# 12648-010), freezing overnight at -80°C in a Mr. Frosty™ Freezing Container (Thermo Scientific™, 5100-001), and storing in a liquid nitrogen dewar. Cryopreserved cells were thawed for culture according to the OncoPro medium user guide.

### Culture of commercially available tumoroid models

Tumoroid models procured from the National Cancer Institute Patient-Derived Models Repository (NCI PDMR; full tumoroid line names and details provided in **Supplementary Table S6**) were cultured in medium outlined in PDMR standard operating procedures, with formulations calling for L-WRN conditioned medium instead using 500 pM Gibco™ Wnt Surrogate-Fc Fusion Recombinant Protein (Thermo Fisher Scientific, cat# PHG0402), 500 ng/mL recombinant RSPO1 (Thermo Fisher Scientific, cat# 120-38-500UG), and 100 ng/ml recombinant noggin (Thermo Fisher Scientific, cat# 120-10C-250UG). The 3dGRO™ human ISO72 colorectal cancer tumoroid line was purchased from Millipore Sigma (Burlington, MA, USA, cat# SCC507) and cultured in vendor-specified homebrew media. Detailed recipes are provided in **Supplementary Table S5**. Embedded cultures were performed via the same methods as tumoroids derived in OncoPro medium, with 50,000 dissociated tumoroid cells embedded in 50 µL Geltrex matrix droplets upon culture initiation and after subculturing. Tumoroids were monitored until tumoroid diameter was approximately 100-300 µm, after which tumoroids were dissociated using Accutase reagent, counted, and reseeded as described above. Where possible, cell pellets for subsequent DNA isolation or cell resuspensions in TRIzol™ reagent were collected as described above at the passages detailed in **Supplementary Table S6**. NCI PDMR tumoroids were also cultured in OncoPro medium in embedded or suspension formats. Tumoroid cells were either seeded in OncoPro medium upon thaw or, in select cases, from active culture in homebrew media (details in **Supplementary Table S6**). For mutational and gene expression analysis, cells from an initial cell bank (either directly as received from vendor or after establishment of an internal cell bank using vendor-recommended culture conditions; see **Supplementary Table S5** and **Supplementary Table S6** for details) were compared to tumoroids expanded from the initial bank in different arms of the study. Some cultures were not processed for RNA sequencing (details in **Supplementary Table S6**).

### 2D cell culture

Colorectal cancer cells RKO (CRL-2577™), SW48 (CCL-231™), SW837 (CCL-235™), SW948 (CCL-237™), and SW116 (CCL-233™) cells were purchased from ATCC (Manassas, VA, USA), thawed, and cultured in Gibco™ DMEM/F-12 (Thermo Fisher Scientific, cat #11320-033), supplemented with 10% Gibco™ fetal bovine serum (FBS), certified, One Shot™ format (Thermo Fisher Scientific, cat #A31604-01) and 1% penicillin/streptomycin (Thermo Fisher Scientific, cat# 15140-148). Cells were cultured in T75 flasks for 72 hours and, after reaching 80-90% confluency, detached using 0.05% Gibco™ Trypsin-EDTA (Thermo Fisher Scientific, cat #25300-054). Cells were counted and preserved for RNA and DNA analysis as described above.

### Targeted genomic sequencing

*DNA extraction, library preparation, and sequencing*: Genomic DNA (gDNA) was isolated from the frozen cell pellets using the Invitrogen™ PureLink™ Genomic DNA Mini Kit (Thermo Fisher Scientific, cat# K182002) following the manufacturer’s protocol. Isolated DNA was quantified using the Invitrogen™ Qubit™ dsDNA High Sensitivity Assay Kit (Thermo Fisher Scientific, cat# Q32854). Sequencing libraries were prepared from 10 ng gDNA using the Ion Torrent™ Oncomine™ Comprehensive Assay v3C (OCAv3C) kit (Thermo Fisher Scientific, cat# A35806) and the Ion Torrent™ Ion Chef™ instrument model 4247 (Thermo Fisher Scientific, cat# 4484177) following manufacturer’s instructions (MAN0015885 (Revision C.0)) using 15 amplification cycles of 8 minutes each. The instrument library preparation protocol pools the 8 barcoded sample libraries together on an equimolar basis with expected total concentration around 100 pM. The concentration of the pooled library was measured using a High Sensitivity DNA ScreenTape (Agilent Technologies, Santa Clara, CA, cat# 5067-5584) and a 4200 TapeStation System bioanalyzer (Agilent Technologies). The pooled library was diluted to 50 pM in Invitrogen™ non-DEPC Treated, Nuclease-Free Water (Thermo Fisher Scientific, cat# AM9938). Libraries were templated and loaded onto Ion Torrent™ Ion 540™ Chips (Thermo Fisher Scientific, cat# A27766) using the Ion Chef™ instrument and the Ion Torrent™ Ion 540™ Kit-Chef (Thermo Fisher Scientific, cat# A30011) according to the manufacturer instructions (MAN0013432, Revision (E.0), MAN0010851 (Revision F.0)). Sequencing was performed using the Ion Torrent™ Ion GeneStudio™ S5 System model 7727 (Thermo Fisher Scientific, cat# A38194) according to the manufacturer instructions.

*QC, filtering, and analysis*: Ion Reporter™ Software v5.18 (Thermo Fisher Scientific) was used to map the data to the human genome assembly version 19 and perform downstream analysis. The Oncomine™ Comprehensive v3-w4.2-DNA-Single Sample analysis workflow was used to align raw reads to the reference genome and identify genomic variants. Coverage analysis reports from the Ion Reporter™ Software providing measurements of mapped reads, mean depth, uniformity, and alignment over a target region were used as quality assessment of the sequencing reactions.

*SNV analysis:* SNV calls with coverage ≥100 were used for correlation analysis. In cases where an SNV was identified in only 1 sample of a comparison, a value of 0 was assigned to that locus in the other sample to represent the reference call (0 frequency for no mutation present/identified at that locus); though in some cases this absence may be due to lack of sequencing depth, we chose the more stringent interpretation by using the 0 value. In cases where multiple samples are cross-correlated at once, the addition of 0 values can increase correlation values between similar samples that both lack mutations at loci present in unrelated samples; this can lead to higher correlation values between samples, as more donors are compared, for example, Fig. 2a colorectal donors D1-D3 versus Fig. 4C but not lung donors Fig. 2a D1 and D2 versus Fig. 4C. The allelic frequency of SNVs at each genomic loci for samples were correlated using Pearson’s correlation in JMP18. For base substitution analysis, the six major single base substitutions: C>A, C>G, C>T, T>A, T>C, T>G were counted using Microsoft Excel. Oncogenic driver mutations were identified using the inbuilt Oncomine™ Variants plugin 5.18 filter in the Ion Reporter™ software. For **Supplementary Fig. S1**, mutations were called using the Oncomine™ Extended filter 5.18 were counted.

*CNV analysis*: Unfiltered results from the Oncomine™ Comprehensive v3-w4.2-DNA-Single Sample analysis workflow were downloaded and sorted for CNV coverage. Estimated ploidy values were cross-correlated by genomic locus in JMP18.

### Tumor mutation load assay

Genomic DNA was isolated and quantified as described above. Sequencing libraries were prepared from 20 ng of gDNA using the Ion Torrent™ Oncomine™ Tumor Mutation Load Assay, Chef-ready kit (Thermo Fisher Scientific, cat# A37910) and the Ion Chef™ following manufacturer’s instructions (MAN0017042 (Revision C.0)) using 13 amplification cycles of 16 minutes each. Library quantification, templating, and sequencing were performed as described above.

### qPCR Genotyping assays

Genomic DNA was isolated and quantified as described above. Purified DNA was amplified using Applied Biosystems™ TaqMan™ Genotyping Master Mix (Thermo Fisher Scientific, cat# 4371357) in the presence of Applied Biosystems™ TaqMan™ SNP Genotyping Assay, human (Thermo Fisher Scientific, cat# 4351379) using an Applied Biosystems™ QuantStudio^TM^ 12K Flex Real-Time PCR System (Thermo Fisher Scientific, cat# 4471087) according to manufacturer’s protocol. Both commercially available and custom designed primer and probe combinations were screened to identify mutations in the *APC* gene (**Supplementary Table S4**).

### Bulk RNA sequencing

*RNA extraction, library preparation, and sequencing*: Cell pellets were dissolved in TRIzol™ reagent and RNA was isolated using the Invitrogen™ PureLink™ RNA Mini Kit (Thermo Fisher Scientific, cat# 12183018A). RNA integrity and concentration were determined using a High Sensitivity RNA Screen Tape (Agilent, cat# 5067-5579) and a 4200 TapeStation System bioanalyzer (Agilent); RNA concentration was quantified using the Invitrogen™ Qubit™ RNA High Sensitivity Assay Kit (Thermo Fisher Scientific, cat# Q32852). DNA digestion was performed using the Invitrogen™ ezDNase™ enzyme (Thermo Fisher Scientific, cat# 11766051). Complementary DNA (cDNA) was then synthesized from 10 ng RNA using the Invitrogen™ SuperScript™ IV VILO™ Master Mix kit (Thermo Fisher Scientific, cat# 11766050) or Invitrogen™ SuperScript™ VILO™ cDNA Synthesis Kit (Thermo Fisher Scientific, cat# 11754050) following the manufacturer protocol. Sequencing libraries were prepared from the total cDNA synthesis product using the Ion Torrent™ Ion Ampliseq™ Transcriptome Human Gene Expression Panel, Chef-Ready Kit (Thermo Fisher Scientific, cat# A31446) and the Ion Chef™ instrument model 4247 following manufacturer’s instructions (MAN0010742 (Revision C.0)) and using 12-13, 16-minute amplification cycles. The pooled library concentration was measured using a High Sensitivity D1000 ScreenTape (Agilent, cat# 5067-5584) and a 4200 TapeStation System bioanalyzer (Agilent). The pooled library was diluted to 70 pM using Invitrogen™ non-DEPC treated, nuclease free water (Thermo Fisher Scientific, cat# AM9939). Templating and sequencing were performed as described above. In occasional cases of low read quality, reads could be increased by heat-denaturing an aliquot of the prepared sequencing library at 99°C for 5 minutes and then placing on ice for 2 minutes immediately before templating (updated guidance from MAN0013432 (Revision K.0)).

*QC, filtering and analysis*: The sequencing data were automatically transferred to the S5 Torrent server virtual machine for alignment, quality control, and analysis using the human genome assembly 19 as the reference genome. Sequence read alignment was performed using the hg19 human reference file: ‘hg19_ampliseq_transcriptome_ercc_v1’. The AmpliseqRNA plugin was used to target regions of the 21K reference sequence genes using the hg19_AmpliSeq_Transcriptome_21K_v1 target panel to generate reads matrix data. Only samples with reads above 5 million were considered for further downstream analysis. Differential gene expression analysis was performed using DESeq2 (version 3.19)^59^ using false discovery rate (FDR)*<*0.05 and log2 fold-change (absolute value *>*1) parameters to define differentially expressed genes (DEGs). DEGs were used for further downstream pathway enrichment analysis, performed using ShinyGO 0.77^60^. Enriched Kyoto Encyclopedia of Genes and Genomes (KEGG) pathways are displayed. In cases where more than 10 pathways were enriched, the top 10 results (ranked by FDR) are shown. FDR>0.05 was used as a threshold to identify significantly enriched KEGG pathways. PCA analysis and sample-to-sample distance analysis were performed using variance stabilizing transformed (VST) data in DESeq2. For sample-to-sample distance analysis, the R function *dist* was used to calculate Euclidean distance between samples. For combined analysis of samples from multiple sequencing runs, gene expression data from all the cancer indications and donors were first normalized for sequencing depth, log_2_ transformed, and mean centered by genes (rows). Genes that had greater than 0.5 reads per million in at least two samples were filtered in to remove lowly expressed genes. The top 50 variable genes from the normalized gene expression data were calculated using the R function *var* and genes were clustered using hierarchical clustering.

### Molecular subtyping of initial cancer cells samples and tumoroids

*Molecular subtyping of colorectal tumoroids*: CRCAssigner, an R software package (https://github.com/syspremed/CRCAssigner), was used to predict consensus colorectal cancer molecular subtypes^19^. The package calculates the Pearson’s correlation between each sample expression profile and 38 marker genes (CRC-38) using the Prediction Analysis of Microarray (PAM) centroid method. Samples are subtyped into five major clinical subtypes, namely stem-like, inflammatory, transit amplifying, goblet-like, and enterocyte, based on the highest correlation. The PAM786 gene list^19^ was used for comparison of colorectal tumoroid lines to colorectal cancer tumor samples and established 2D colorectal cancer cell lines.

*Molecular subtyping of breast tumoroids*: PAM50, an R software package (https://github.com/ccchang0111/PAM50), was used to classify breast cancer samples into 5 major subtypes: Normal, Luminal B, Luminal A, Her2, and Basal-like based on gene expression levels of the PAM50 genes.

### Single-cell RNA sequencing

*Library preparation and sequencing*: Cryopreserved single cells from dissociated colorectal tumors and tumoroids were submitted to Azenta Life Sciences (Burlington, MA, USA) for scRNA-seq. The sample libraries were prepared in a single batch using the Chromium Single-Cell 3′ v3 Library and Gel Bead Kit (10x Genomics). Briefly, gel bead-based emulsions (GEM) were generated by combining barcoded single-cell 3′ Gel Beads, cells, and partitioning oil. Ten times barcoded, full-length cDNAs generated from GEMs were amplified by polymerase chain reaction (PCR). Enriched libraries were enzymatically digested, size selected, and adaptor ligated for sequencing. Sequencing libraries were generated with unique sample indices for each sample and quantified using the Kapa library kit. Quantified libraries were sequenced on an Illumina NovaSeq™ 6000.

*QC, filtering and analysis*: cellranger-7.1.0 Single-Cell Software Suite from 10x Genomics (https://www.10xgenomics.com/support/software/cell-ranger/latest) was used to align FASTQ sequencing reads to the cellRanger_GRCh38_v5 reference transcriptome, generating single-cell feature counts and associated unique molecular identifiers (UMIs) for each sample. The Seruat R package (version 4.3.0) was used to pre-process single-cell data. The CreateSeuratObject function was used to create Seurat objects for the count data. All the 10x genomics Seurat objects were merged using the merge function in Seurat 4.3.0. Genes/features shared by three or more cells, cells that expressed ≥200 and ≤5,000 genes, and mitochondrial gene content <20% were included for analysis. The gene expression measurements for each cell were normalized by the total expression, multiplied by a scale factor of 10,000, and then log transformed. The FindVariableFeatures function was used to identify the top 2,000 highly variable genes. To scale the expression of each gene, data was transformed linearly so that the mean was 0 and variance was 1. PCA was performed on the scaled 2000 highly variable genes. Data were visualized using Uniform Manifold Approximation and Projection (UMAP). Cell clusters were identified by a shared nearest-neighbor (SNN) modularity optimization-based clustering algorithm set at a resolution of 0.5. To identify differentially expressed genes for cluster demarcation, the FindAllMarkers module was used, and genes expressed in more than 25% of the cells in each cluster were selected. The top 20 differentially expressed genes markers from each cluster were used to annotate cell types.

Gene expression data of 18,042 single cells from two colorectal tumors and tumoroids were aggregated first, and, using unsupervised clustering, cells were then clustered together in two-dimensional space. Epithelial cell adhesion molecule (*EPCAM*) was used to identify epithelial cell clusters. Collagen 1 A1 (*COL1A1*) was used to identify fibroblast cell clusters, and the mesenchymal marker vimentin (*VIM*) was used to identify stromal and immune cell clusters. Protein tyrosine phosphatase receptor type C (*PTPRC*) or leucocyte common antigen (*CD45*) was used to distinguish immune cells from stromal cells (**Supplementary Fig. S4**).

Raw gene read counts with pre-annotated cell types (immune cells, epithelial cells, endothelial cells, fibroblast cells) were transcript per million (TPM) normalized and log2 transformed for dissociated tumor cells (P0), tumoroids at passage 10 (P10) and tumoroids at passage 27 (P27) for HuCo3209 and HuCo021320 colorectal donors. In each donor, the genes across all the valid cells across passages with mean(log2(TPM))>=1.5 were selected to construct donor gene expression matrix *M* (*donor*)=genes×cells across different passages (P0, P10, and P27). Genes with enough reads were retained as initial input to infer Copy Number Variation (CNV) and were ordered by genomic coordinates from chr1 to chr22. Immune cells from the P0 cells were used as the “referenced group”, to infer CNV from EPCAM positive cells in matrix *M* (donor) as the “observed group”. Using R package inferCNV (https://github.com/broadinstitute/inferCNV) with default parameters compatible to 10x genomics scRNA-seq library, the CNV profile for each cell was estimated.

### In vivo tumorigenicity experiments and histology

Tumoroids were expanded in suspension culture in OncoPro medium per manufacturer’s instructions before being dissociated, counted, and resuspended in DPBS(+/+)containing 33% Corning^®^ Matrigel^®^ basement membrane extract (BME) such that a 100 µL injection volume contained 1e6, 3e6, or 5e6 cells. Cells were injected at 1e6, 3e6, or 5e6 cells per NSG mouse subcutaneously, with three animals per condition, at the University of Maryland Greenebaum Comprehensive Cancer Center Translational Laboratory Shared Service (TLSS) facility. All experiments were performed under IACUC approval by the University of Maryland Baltimore Animal Care and Use Program. Tumor volume was monitored for up to 90 days post-injection, at which time mice were euthanized by CO_2_ inhalation. Mice were euthanized prior to this time point if tumors reached 10% of mouse body weight, or if no tumor was present at 60 days post-injection. Tumor formation was most robust at 3e6 cells per mouse injected, and data from that condition was used for **Supplementary Fig. S6**. Following euthanasia, tumors were excised and placed in fixative. Tumoroids from matched lines were cultured in suspension using OncoPro medium, collected, washed in cold DPBS (-/-), and fixed for 1 hour at room temperature in Invitrogen™ Image-iT™ Fixative Solution (Thermo Fisher Scientific, cat# FB002). Fixed tumor samples and tumoroids were washed twice with DPBS (-/-) and then dehydrated by sequential washes with 30% ethanol, 50% ethanol, and 70% ethanol solutions (all diluted in water). Samples were stored in 70% ethanol solution prior to being processed for histology. Samples were paraffin embedded, sectioned, affixed to slides, and stained by Histoserv, Inc. (Rockville, MD, USA). Slides were stained with H&E, Jone’s PAMS, Movat Pentachrome, and Dane’s Method stains. Images of stained slides were acquired on an Invitrogen™ EVOS™ M7000 Imaging System.

### Generation of stable GFP-expressing tumoroid line

A transfer plasmid carrying an EmGFP gene under the EF1a promoter and a blasticidin resistance gene was transfected into human embryonic kidney (HEK) 293-derived Gibco™ Viral Production Cells using the Gibco™ LV-MAX™ Transfection Kit (Thermo Fisher Scientific, cat# A35684) to produce the lentivirus (LV) required for transduction. Prior to the transduction, HuCo1044 tumoroids were dissociated into a single-cell suspension. The number of viable cells was determined using the Invitrogen™ Countess™ 3 FL Automated Cell Counter (Thermo Fisher Scientific, cat# AMQAX2000). Cells were seeded into a 6-well plate at a seeding density of 2e5 viable cells per mL of OncoPro medium with a final volume of 2 mL per well. Transduction was performed in supplemented OncoPro medium at a multiplicity of infection (MOI) of 1, 5, 10, 25, and 50 in order to determine the optimal level. An MOI of 50 yielded the highest percent of GFP-positive cells as measured using an Invitrogen™ Attune™ NxT Flow Cytometer (Thermo Fisher Scientific, cat# A24858), and this sample was used for the rest of this study. Twenty-four hours post-transduction, cells were reseeded into standard suspension culture conditions in selection medium containing 5 µg/mL blasticidin. Cells were passaged every 7–14 days depending on tumoroid size, with media changes performed every 2–3 days. Four weeks after selection began, the cells were recharacterized by flow cytometry for GFP expression and by next generation sequencing using the Ion Torrent™ Oncomine™ Comprehensive Assay v3C and Ion Torrent™ Ion Ampliseq™ Transcriptome Human Gene Expression Panel as described above.

### Flow cytometry for phenotyping of dissociated tumor cells and established tumoroids

Tumor samples or tumoroid cultures were dissociated as described above and passed through a 100 µm strainer. Briefly, cells were resuspended to 1e6 cells/mL in DPBS(+/+), and 1e4 to 1e5 cells were plated per well of a U-bottom 96-well plate. Cells were incubated with 1:500 Invitrogen^TM^ LIVE/DEAD^TM^ Fixable Yellow Dead Cell Stain (Thermo Fisher Scientific, cat# L34959) for 30 minutes at 4°C and then washed with Invitrogen^TM^ eBioscience^TM^ Flow Cytometry Staining Buffer (Thermo Fisher Scientific, cat# 00-4222-57). Cells were incubated with Invitrogen™ Fc Receptor Binding Inhibitor Polycolonal Antibody, eBioscience™ (FcBlock; Thermo Fisher Scientific, cat# 14-9161-73) at 20% concentration in Flow Cytometry Staining Buffer at 4°C for 20 minutes. Primary antibodies against surface targets were added and incubated for 1 hour at 4°C. Antibodies were directed against EpCAM (1:20 final dilution; Thermo Fisher Scientific, cat# 25-9326-42, PE-Cyanine7), CD31 (1:20 final dilution; Thermo Fisher Scientific, cat# 46-0319-42, PerCP-eFluor™), CD45 (1:10 final dilution; Thermo Fisher Scientific, cat# 12-0459-42, PE), and/or CEACAM (1:20 final dilution; Thermo Fisher Scientific, cat# 62-0668-42, Super Bright™ 436). Samples were washed three times with Flow Cytometry Staining Buffer and then fixed and permeabilized using the Invitrogen^TM^ eBioscience^TM^ Foxp3/Transcription Factor Fixation/Permeabilization Concentrated and Diluent (Thermo Fisher Scientific, cat# 00-5521-00) working solution according to the manufacturer’s protocol. Samples were blocked in 10% Gibco™ goat serum, New Zealand origin (Thermo Fisher Scientific, cat# 16210-064) in 1X Invitrogen™ eBioscience™ Permeabilization Buffer (Thermo Fisher Scientific, cat# 00-8333) for 15 minutes at room temperature. Intracellular antibody (against vimentin; 1:50 final dilution; Thermo Fisher Scientific cat# MA5-28601, APC) was added and incubated for 1 hour at room temperature. Samples were washed twice and then resuspended in 200 µL Flow Cytometry Staining Buffer and analyzed on the Attune Nxt Flow Cytometer using the Invitrogen™ CytKick^TM^ Autosampler (Thermo Fisher Scientific, cat#, A42901). Bead-based compensation (Invitrogen™ UltraComp eBeads™ Plus Compensation Beads, Thermo Fisher Scientific, cat# 01-3333-42 and Invitrogen™ ArC™ Amine Reactive Compensation Bead Kit, Thermo Fisher Scientific, cat# A10346) was applied and fluorescence minus one controls were used to set gates for expression.

### Preparation of the effector cells for NK-tumoroid co-culture

The natural killer (NK) cell line, NK-92, was procured from ATCC (CRL-2407™) and cultured in Gibco™ RPMI 1640 Medium, GlutaMAX^TM^ Supplement (Thermo Fisher Scientific, cat# 61870-036) + 10% Gibco™ fetal bovine serum, qualified, heat inactivated (Thermo Fisher Scientific, cat# 16140-071) + 10% Gibco™ horse serum, heat inactivated, New Zealand origin (Thermo Fisher Scientific, cat# 26050-088) + 500 U/mL Gibco™ recombinant human (rh) IL-2 (Thermo Fisher Scientific, cat# PHC0021). For primary NK cells, human leukopaks were procured from the San Diego Blood Bank and processed using the Gibco™ CTS™ Rotea™ Counterflow Centrifugation System (Thermo Fisher Scientific, cat# A50757/A50760). Primary NK cells were enriched by negative selection using the Invitrogen™ Dynabeads™ Untouched™ Human NK Cells Kit (Thermo Fisher Scientific, cat# 11349D) and expanded in Gibco™ CTS™ NK-Xpander™ Medium (Gibco™, A50190-01) supplemented with 5% hAB serum (Fisher Scientific, cat# BP2525-100) and 500 U/mL rhIL-2, all according to manufacturer instructions. Primary NK cells were characterized via flow cytometry prior to initiation of the co-culture experiment.

### NK-tumoroid co-culture

Dissociated HuCo1044-GFP reporter tumoroid cells were seeded into a black-walled microwell plate with 400 µm diameter microcavities (Gri3D^®^, Sun Bioscience, Lausanne, CH). Cells were seeded at 3×10^5^ viable cells per mL of supplemented OncoPro medium, 50 µL per well, and allowed to settle for one hour at 37°C, 5% CO_2_. After the one-hour incubation, supplemented OncoPro medium + 2.67% v/v Geltrex matrix was added to each well to reach a final Geltrex concentration of 2% (v/v) and 10 µM Y27632. The cells were incubated and monitored for tumoroid formation for 3 days using an Incucyte^®^ SX5 Live-Cell Analysis System (Sartorius, Goettingen, DE, cat# 4816). On day 4, the effector cells (NK-92 or primary NK) were prepared for addition to the tumoroid culture at a variety of effector to target (E:T) ratios (0:1, 0.625:1, 1.25:1, 2.5:1, 5:1, and 1:0), wherein the number of target cells was estimated based on the known doubling time of the tumoroid line. The NK cells were resuspended so that the appropriate number of cells for a given E:T was contained in 50 µL to be added to each well, and cells were resuspended in supplemented NK-Xpander medium plus 500 U/mL rhIL-2 for addition. Minimizing disturbance of intact tumoroids, as much medium as possible was removed from each well of the microwell plate. Effector cells were then added in 50 µL on top of the tumoroids. The plate was incubated for 1 hour at 37°C, 5% CO_2_ to allow the NK cells to settle. After incubation, an additional 150 µL of a 1:1 mixture of NK-Xpander complete medium:OncoPro complete medium containing 500 U/mL rhIL-2 and 10 µM Invitrogen™ CellEvent™ Caspase-3/7 Red Detection Reagent (Thermo Fisher Scientific, cat# C10430) was added to each well. No Geltrex or Y27632 was added at the time of NK cell addition on day 4. The cells were incubated and monitored for GFP and Caspase-3/7 Red reagent signal for 3 days post-initiation of co-culture on the Incucyte platform.

## Acknowledgements

Tumoroid models were provided by the NCI PDMR^20^ where indicated and were developed by the NCI PDMR or their Contributing Institutions (Supplementary Table S6). These models are available from the NCI PDMR. Full details are provided in Supplementary Table S6. We thank the PDMR and their Contributing Institutions for their contributions to this work. We also thank all tissue donors who provided material used in this study.

## Author contributions

CDP, CY, PST, BB, SS, AB, SB, AC, JY, GW, ID, SH, AD, LBS, LZ, VC, JN performed experiments and analyzed data. CDP, CY, and PST prepared figures. CDP, CY, PST, BB, and MD wrote the main manuscript text. All authors reviewed the manuscript.

