## Supplementary Figures for "Long-term maintenance of patient-specific characteristics in tumoroids from six cancer indications in a common base culture media system"

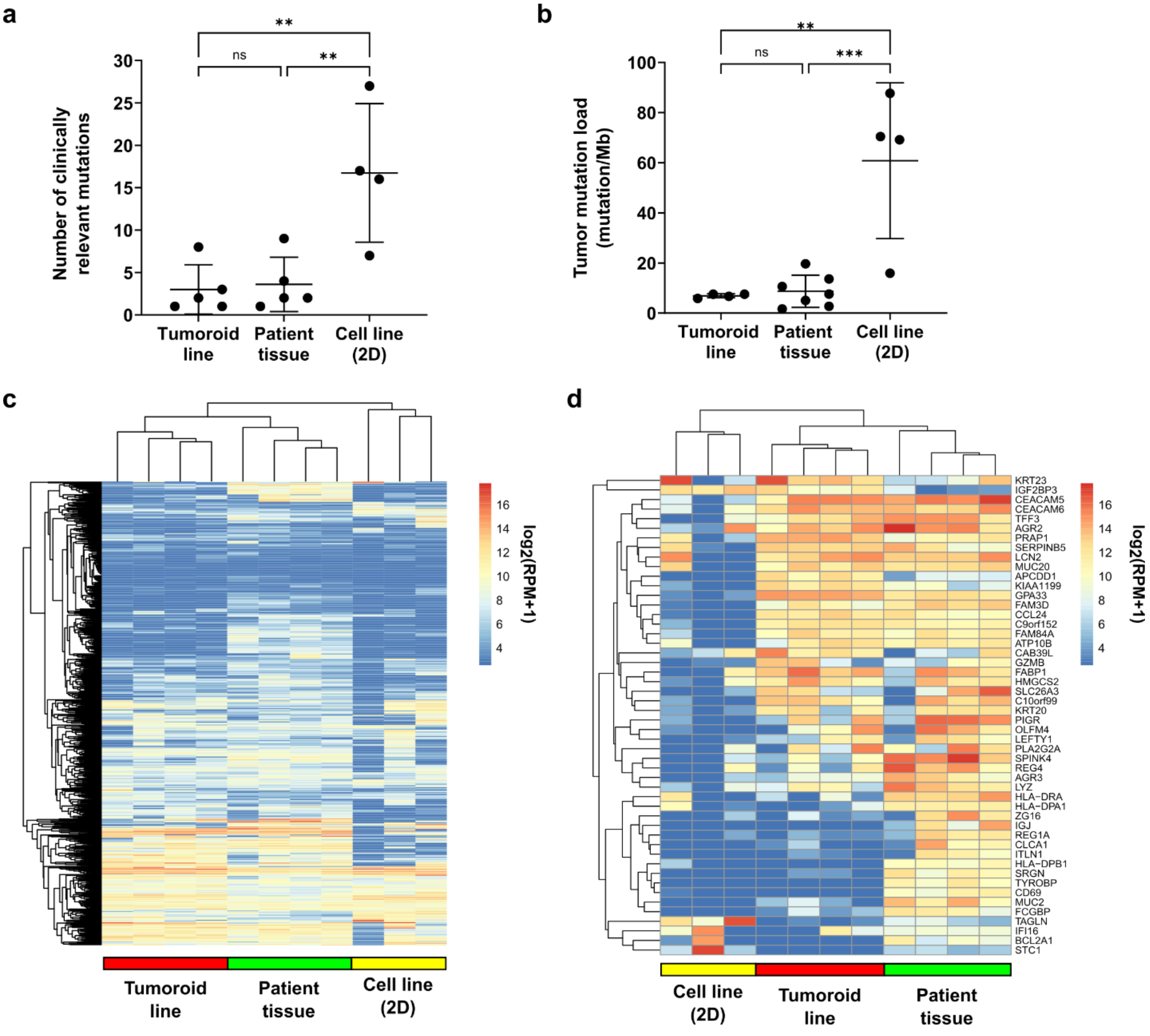


**Supplementary Fig. S1. Comparison of two-dimensional (2D) colorectal cancer cell lines to colorectal tumoroids and colorectal patient tissue samples.** (a) Number of clinically relevant oncogenic mutations present in unmatched colorectal tumoroids, colorectal tumor tissue samples, and 2D colorectal cancer cell lines. The Oncomine™ Comprehensive Assay V3 (OCAv3) was used to capture clinically relevant mutations, which were called using the Oncomine™ Extended filter 5.18 within the Ion Reporter™ software. (b) Tumor mutation load (mutation/Mb) in multiple colorectal tumoroid lines, colorectal tumor tissues, and conventional 2D lines. The Oncomine™ Tumor Mutation Load Assay (TML) was used to quantify tumor mutation load in different samples. Multiple comparison testing (one-way ANOVA followed by Tukey's multiple comparisons test) was performed to compare differences across the three samples. ** and *** represent statistical significance at p-value<0.01 and p-value<0.001, respectively, and ns represents no statistical difference. (c) Gene expression analysis of 751 PAM786 genes in tumor tissue (n=4), tumoroid lines (n=4), and 2D lines (n=4). (d) Heatmap of the top 50 variable PAM786 genes. Rows and columns were clustered using hierarchical clustering. Bulk RNA sequencing was performed using the Ion Ampliseq™ Transcriptome Human Gene Expression Kit targeting >20,000 RefSeq genes.

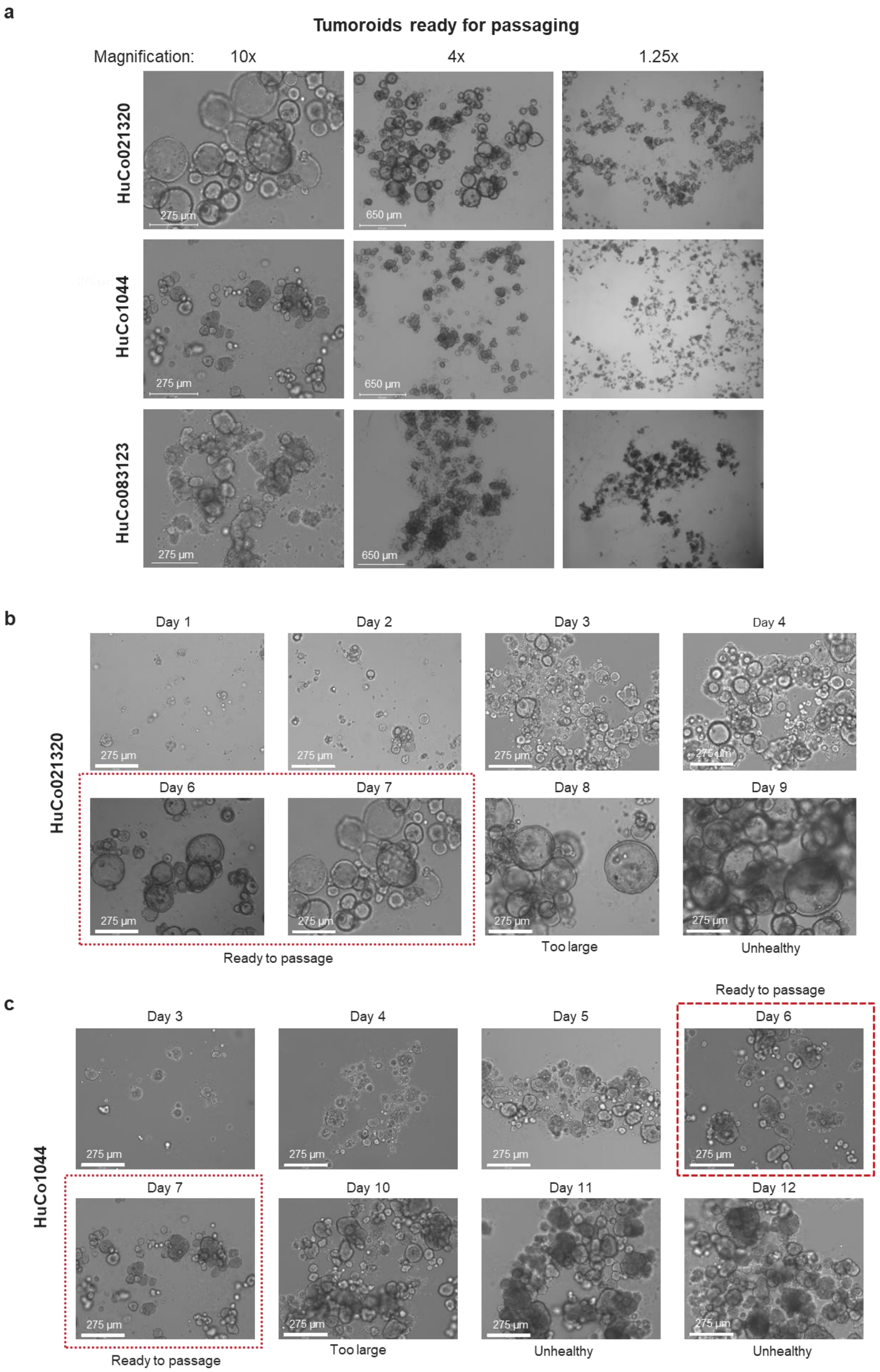


**Supplementary Figure S2. Time course images of tumoroid growth in OncoPro medium.** Tumoroid cells cultured in OncoPro medium in a suspension format were imaged at various magnifications when ready to passage. Three human colorectal tumoroid lines (HuCo021320, HuCo1044, and HuCo083123) are shown. Time course images (10x magnification) of tumoroids during suspension culture in OncoPro medium for (b) HuCo021320 and (c) HuCo1044 tumoroids. Tumoroids that are ready to passage are indicated, as well as overgrown tumoroids, where large size can modulate ability to dissociate tumoroids and/or lead to poor cell health.

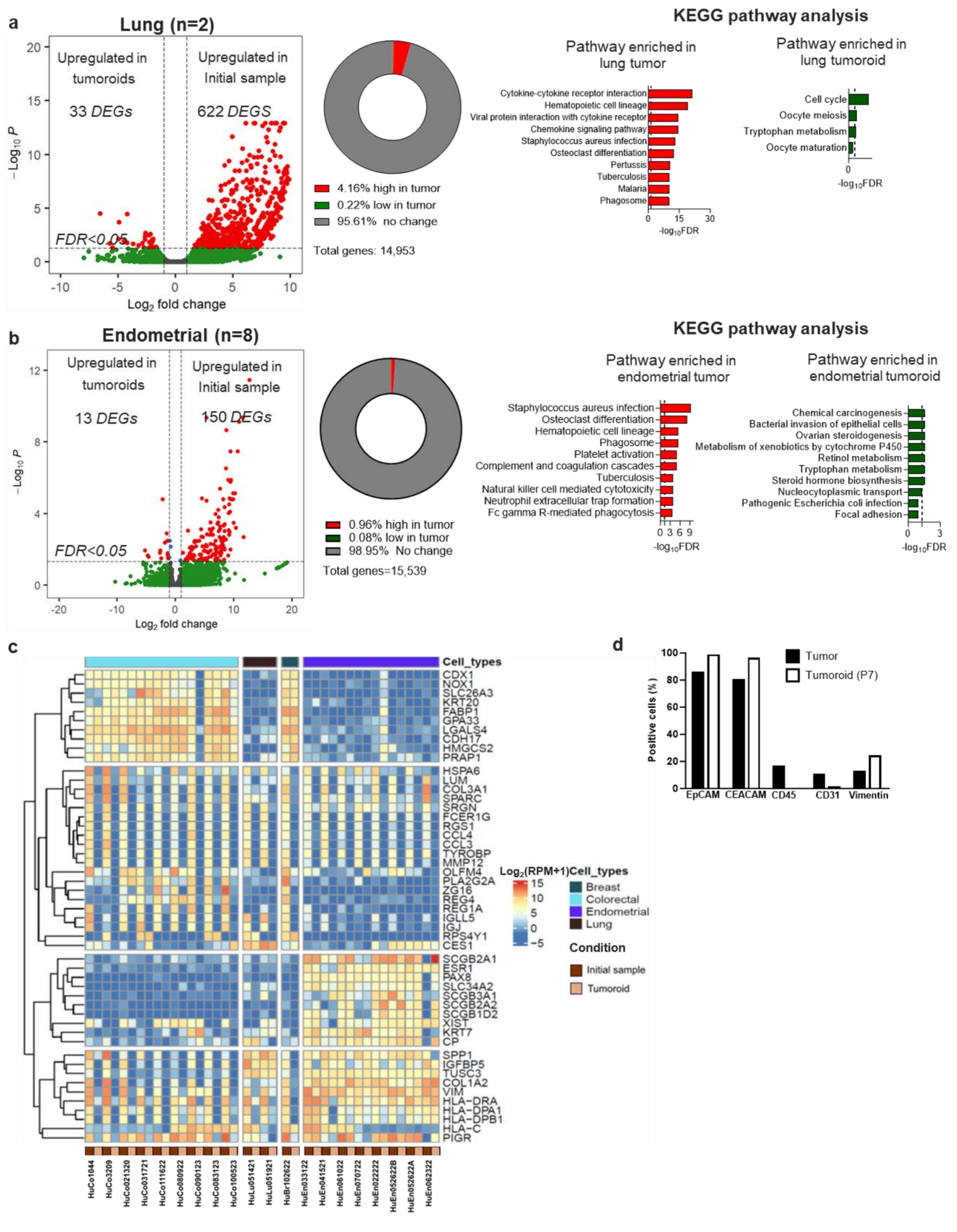


**Supplementary Figure S3.** Differential gene expression analysis of matched tumor tissue and tumoroids in (**a**) lung and (**b**) endometrial cancers at fold change>2 and false discovery rate (FDR)<0.05, and Kyoto Encyclopedia of Genes and Genomes (KEGG) pathway analysis of genes enriched in tumor tissue (top 10 pathways shown, in red) and enriched in tumoroids (top 10 pathways for endometrial shown, in green; all pathways enriched in lung tumoroids are shown). Dotted line indicates FDR=0.05. **(c)** Heatmap of top 50 variable genes obtained from matched tumor tissues and derived tumoroids from colorectal (n=9), lung (n=2), breast (n=1) and endometrial (n=8) cancers. (**d**) Flow cytometry data for expression of epithelial, immune, and stromal cell markers from matched colorectal tumor and tumoroid sample HuCo031721.

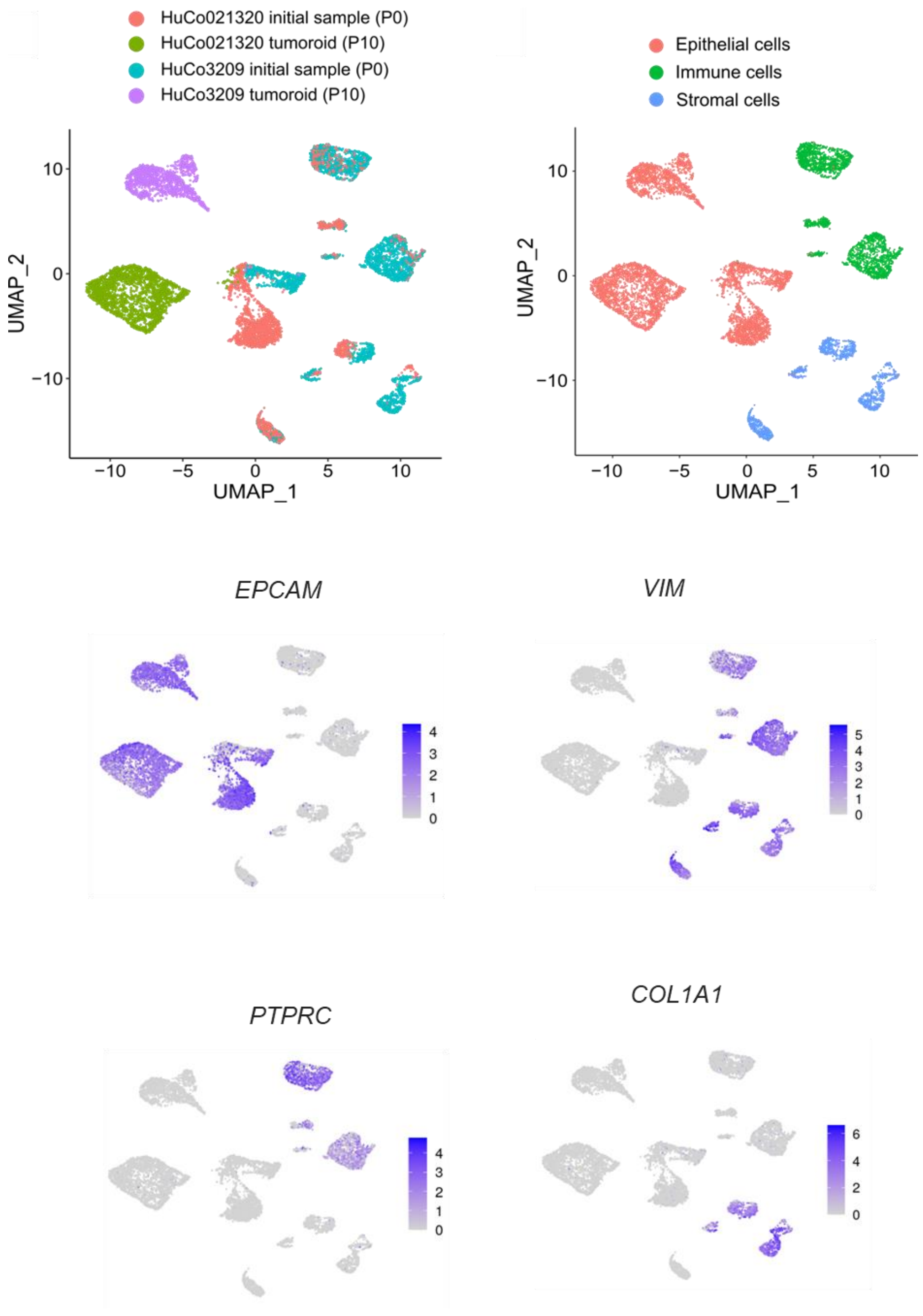


**Supplementary Figure S4**: Feature plot of key gene markers used to identify epithelial cells (epithelial cell adhesion marker, *EPCAM*), mesenchymal cells (vimentin, *VIM*), immune cells (protein tyrosine phosphatase receptor type C, *PTPRC*) and stromal cells (collagen 1 A1, *COL1A1*). UMAP plots in first row are reproduced from Figure 3 for reference.

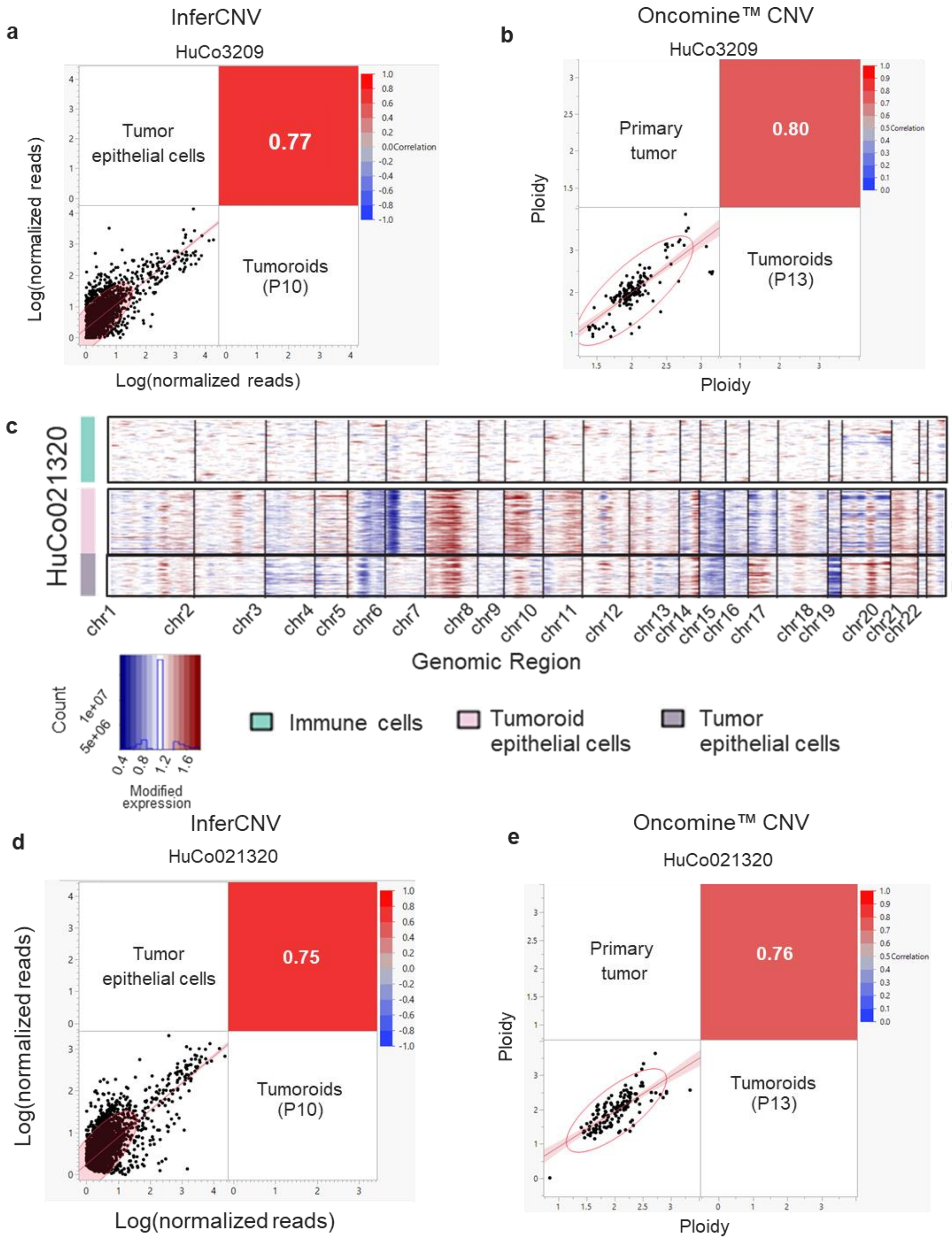


**Supplementary Figure S5. Tumor gene expression and copy number profile is highly concordant with that of derived tumoroids.** (**a**) Pearson's correlation plot of pseudo-bulked single-cell gene expression levels of genes used for inferCNV analysis (n=4,195 genes) from passage 0 (P0) epithelial cells and tumoroids for colorectal model HuCo3209. (**b**) Pearson's correlation plot of ploidy values of primary tumor and tumoroids (HuCo3209) estimated by targeted genomic sequencing with the Oncomine™ Comprehensive Assay v3. Each point represents the ploidy estimate for a given gene. (**c**) InferCNV analysis of P0 HuCo021320 epithelial cells and passage 10 (P10) tumoroid cells, using P0 immune cells as a reference. (**d**) Pearson's correlation plot of pseudo-bulked single-cell gene expression levels of genes used for inferCNV analysis (n=4,973 genes) from P0 epithelial cells and tumoroids for colorectal model HuCo021320. (**e**) Pearson's correlation plot of ploidy values of primary tumor and tumoroids (HuCo021320) estimated by targeted genomic sequencing with the Oncomine™ Comprehensive Assay v3. Each point represents the ploidy estimate for a given gene.

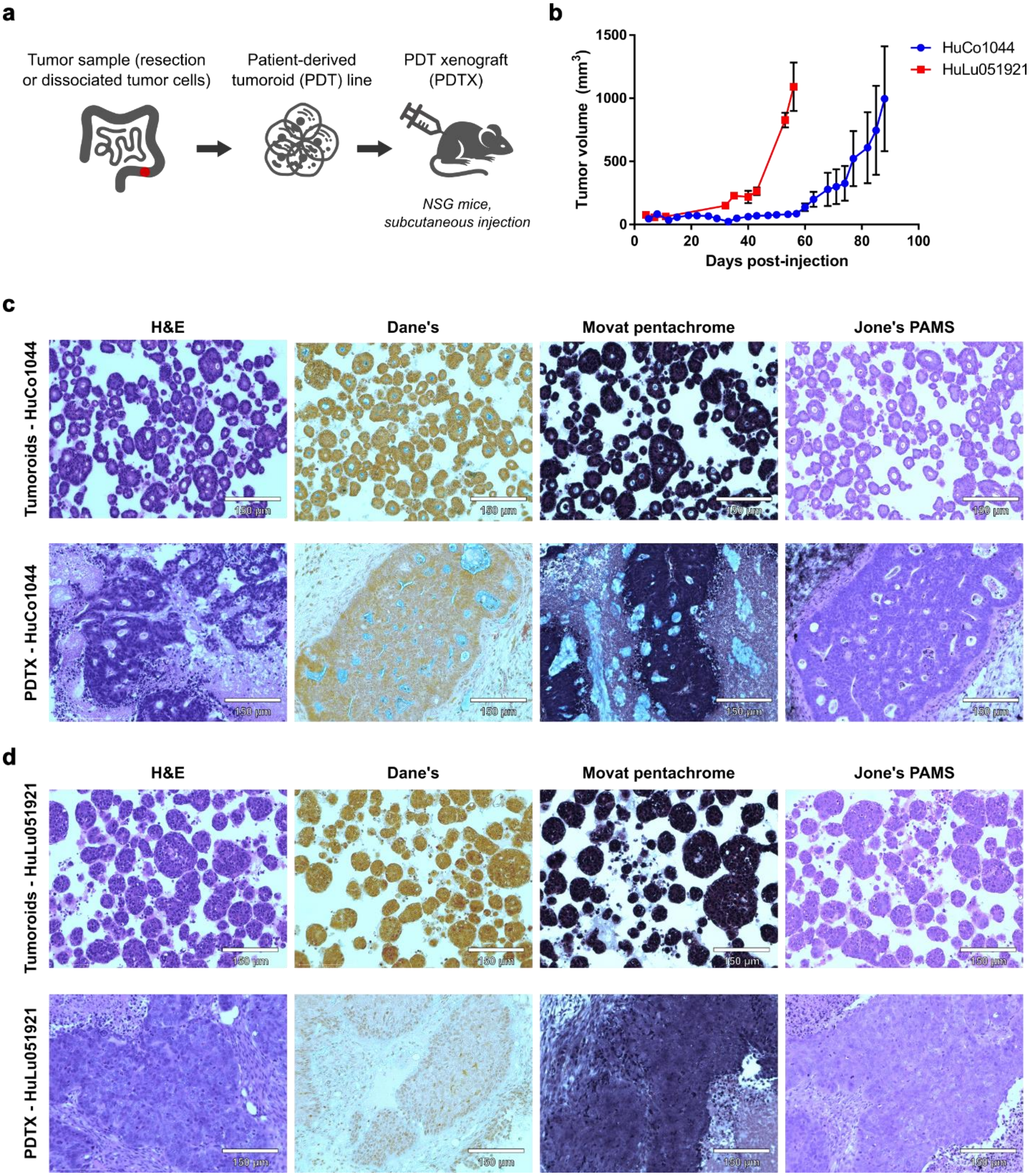


**Supplementary Figure S6**. **Tumorigenicity of established colorectal and lung tumoroid lines.** (**a**) Patient-derived tumoroids were established in OncoPro medium and expanded in suspension culture. Dissociated human colorectal tumoroid cells (HuCo1044) or lung tumoroid cells (HuLu051921) were injected subcutaneously in immunodeficient NSG mice. (**b**) Tumor volume was monitored over time. Plot shows mean ± SEM of tumor volume for 1-3 measurements per time point. Excised tumors or line-matched tumoroids were fixed, embedded, sectioned, and stained with H&E, Dane's, Movat pentachrome, or Jone's PAMS stains for (**c**) HuCo1044 and (**d**) HuLu051921 models.

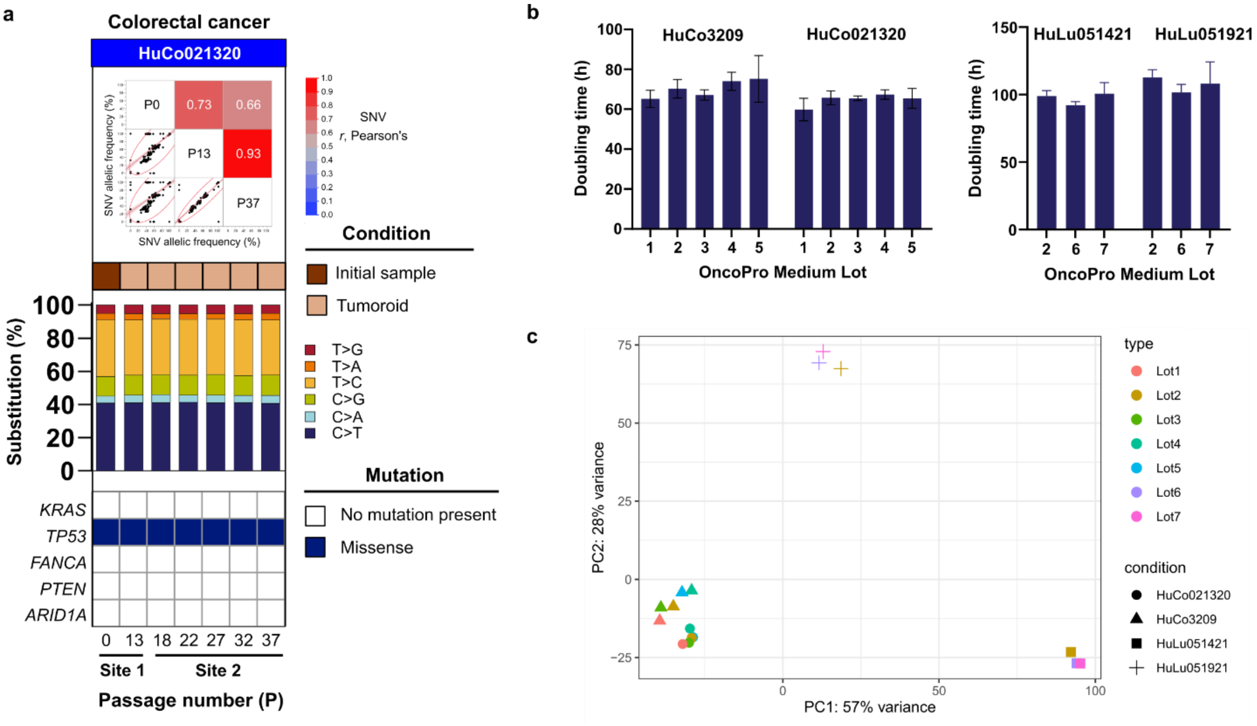


**Supplementary Figure S7. Site-to-site and lot-to-lot consistency using OncoPro medium.** (**a**) A colorectal tumoroid sample, HuCo021320, was established at Site 1 and cryopreserved. Cells from this cryobank were expanded to P50 at Site 1 (see **Figure 4**) and expanded to P37 at Site 2. Comparable genomic stability was observed at both sites based on correlation of SNV allelic frequency (**top**), preservation of single nucleotide substitution frequencies (**bar graphs, middle**), and maintenance of key driver mutations (**heat map, bottom**). In top panel, each dot represents the variant allelic frequency (VAF) for 1 genetic locus covered by the Oncomine™ Comprehensive Assay v3 (OCAv3). The oncogenic mutations were called using the Oncomine™ Variants filter 5.18 within the Ion Reporter™ software. (**b**) Colorectal and lung tumoroids from multiple donors grew at consistent doubling times across multiple lots of OncoPro medium. (**c**) Transcriptomic profiling was performed after culture in each lot of medium. Principal component analysis (PCA) of gene expression profiles demonstrates that tumoroids cluster by sample, rather than medium lot. This testing was performed as part of the long-term culture testing presented in **Figure 5,** and relevant data points from **Figure 5a** are reproduced here, labelled by lot number instead of passage number.

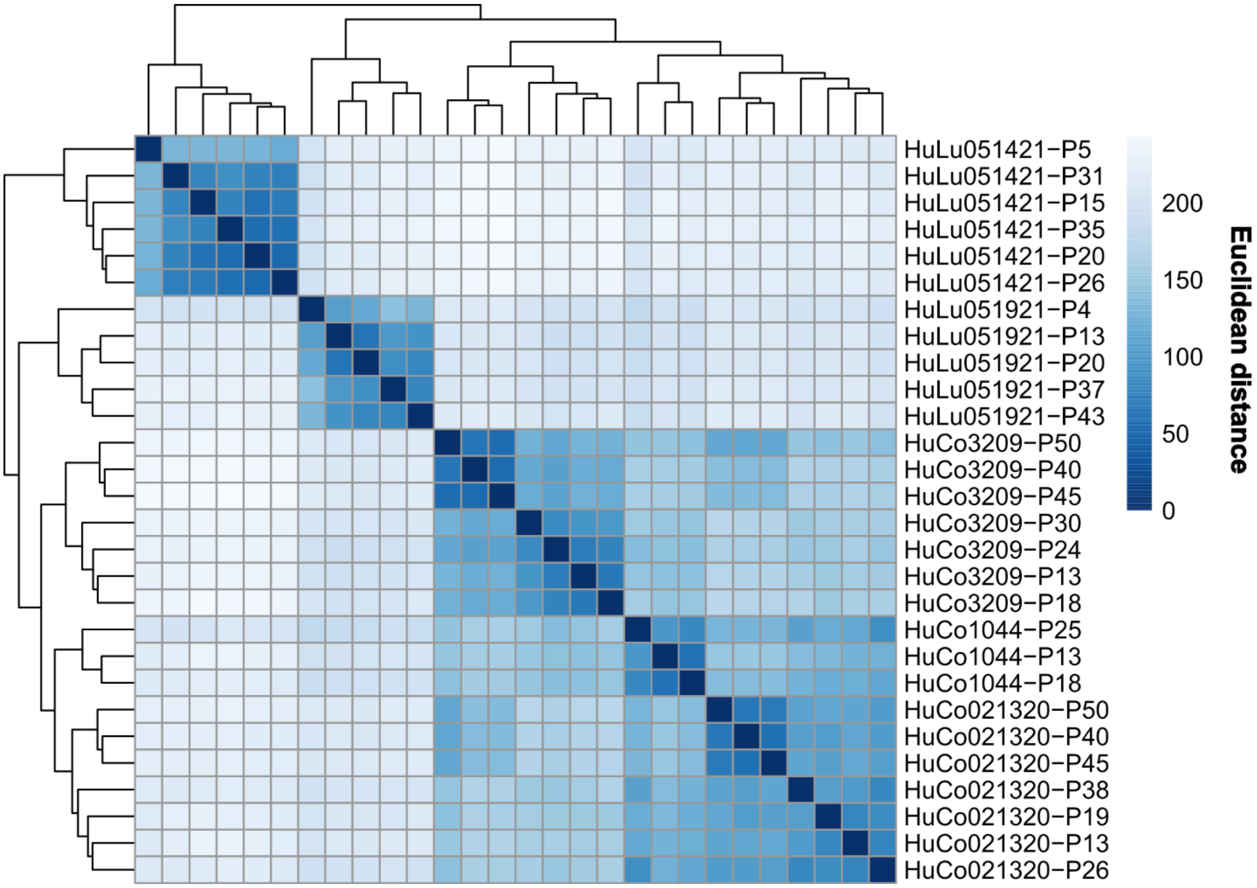


**Supplementary Figure S8. Unsupervised hierarchical clustering of sample-sample Euclidean distance based on gene expression levels in early passage (P) and late passage tumoroids.** Early and late passage tumoroids within a given established human colorectal (HuCo) or lung (HuLu) tumoroid line cluster together.

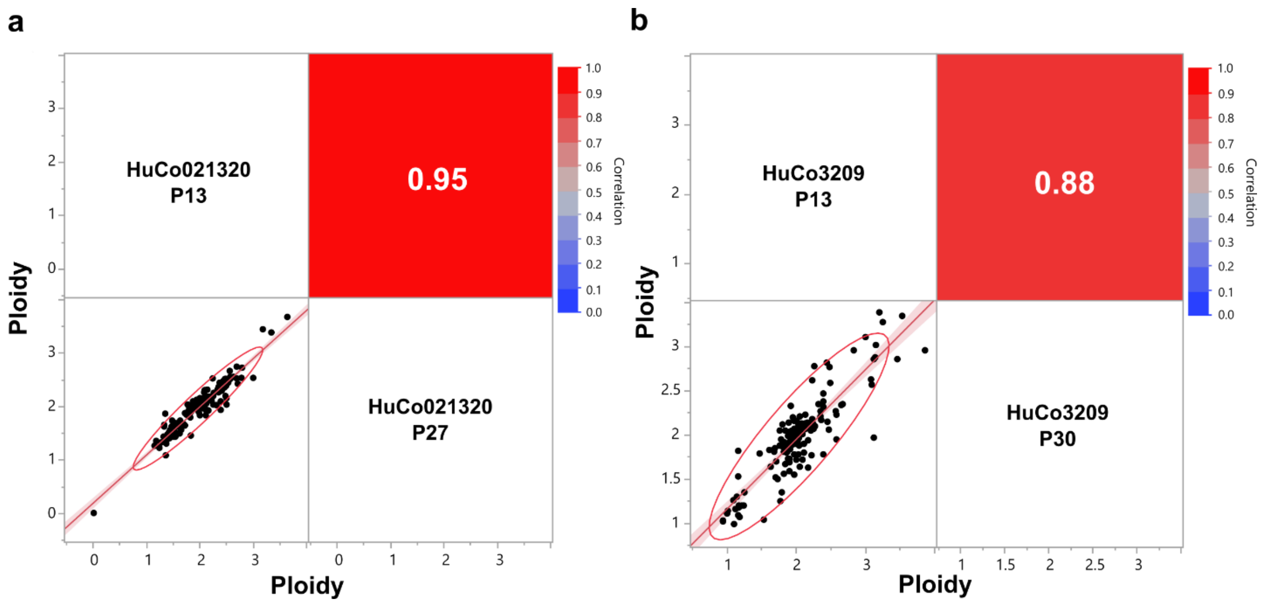


**Supplementary Figure S9.** Correlation of copy number variations (CNV) estimated by targeted genomic sequencing with the Oncomine™ Comprehensive Assay v3, which are maintained during culture of established **(a)** HuCo021320 and **(b)** HuCo3209 tumoroids. Each point represents the ploidy estimate for a given gene.

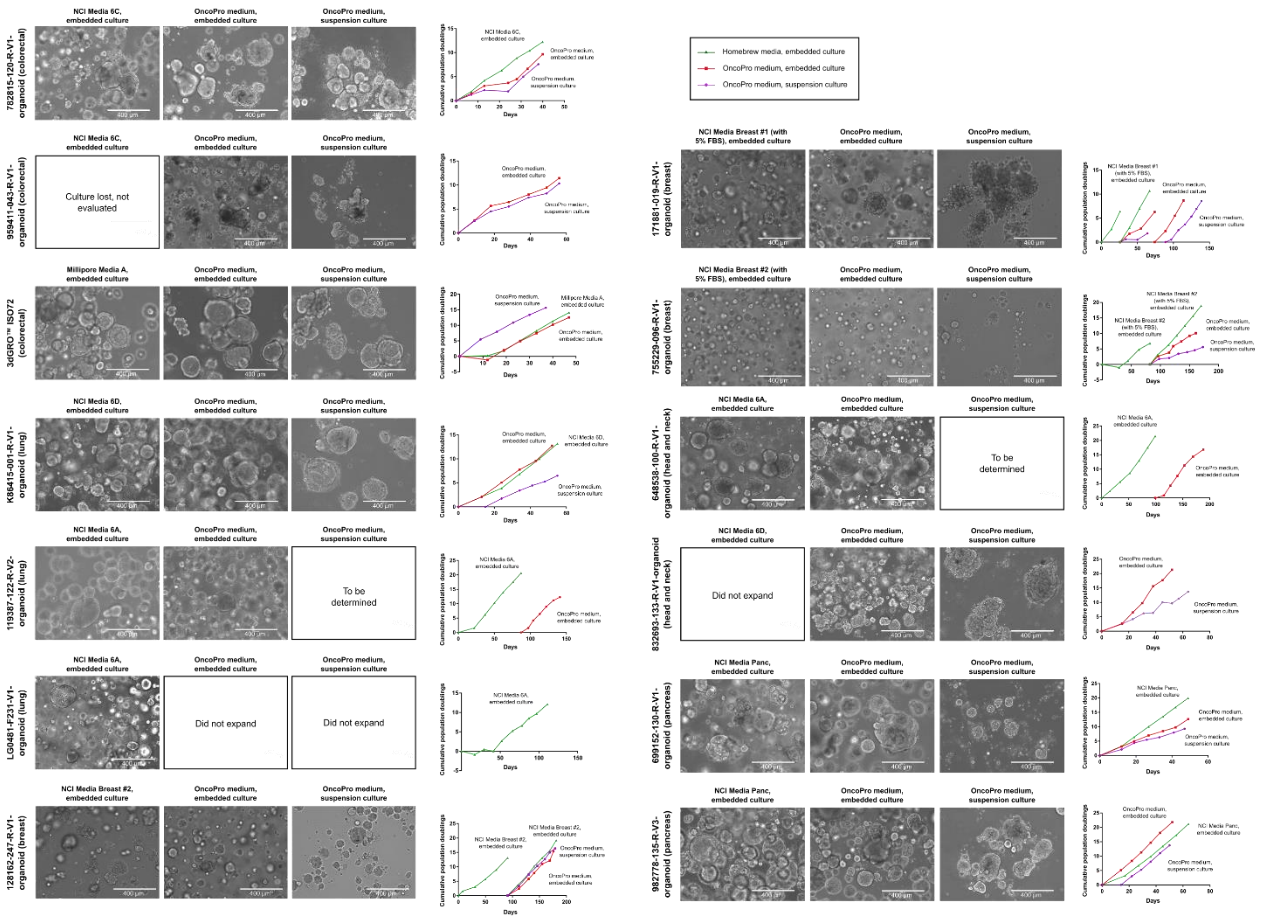

**Supplementary Figure S10. Culture of established, publicly available tumoroid lines in OncoPro medium.** Representative images and growth rates (cumulative population doublings over time) of tumoroid lines across various cancer indications cultured in homebrew media (embedded culture) or OncoPro medium in embedded or suspension culture formats. Scale bar = 400 µm.

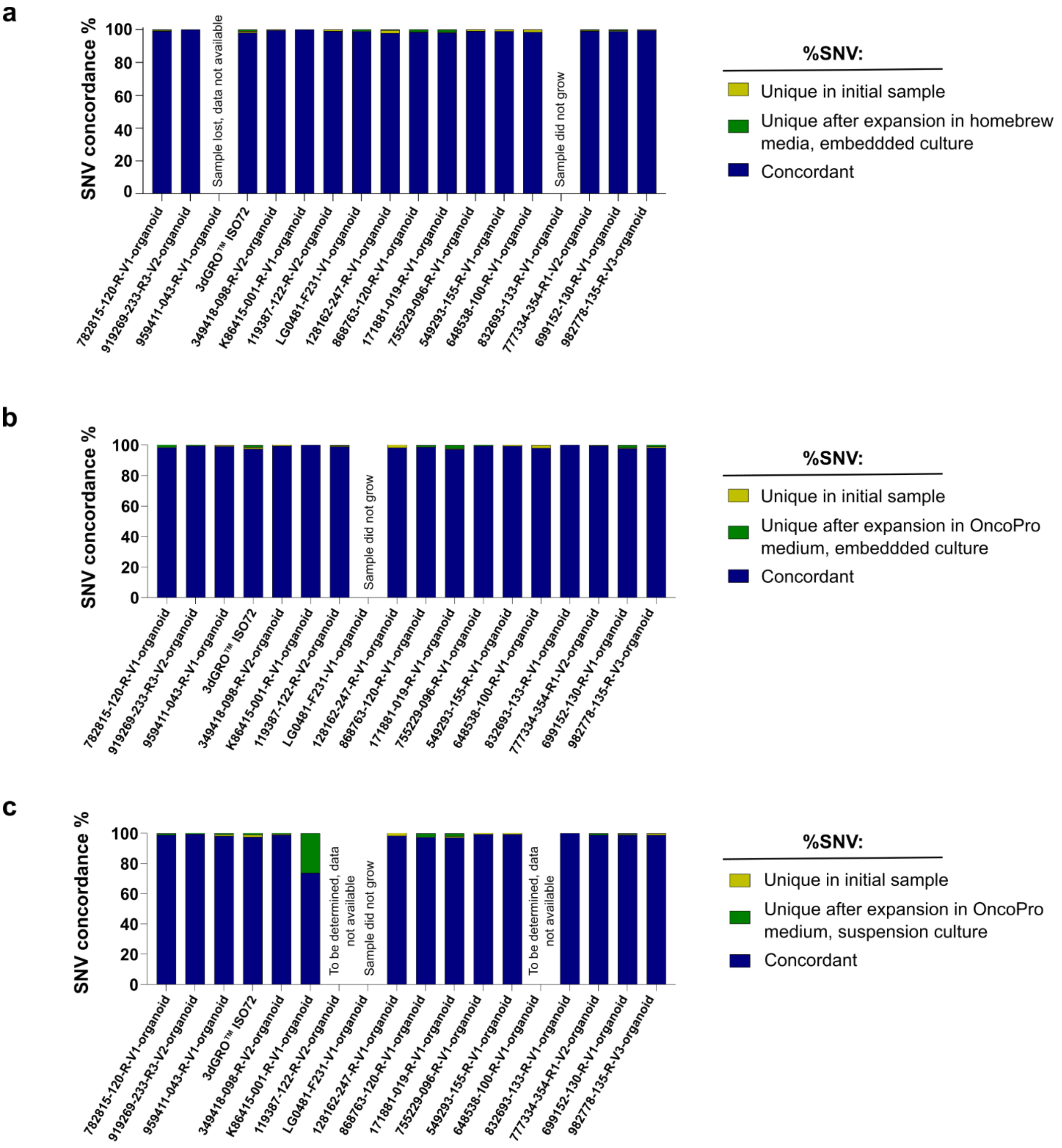


**Supplementary Figure S11. Mutations in tumoroids cultured in homebrew media and in OncoPro medium (embedded and suspension culture) are highly concordant with the initial starting tumoroid material.** (a) Percentage overlap of single nucleotide variants called in initial tumoroid sample (or starting cell bank) and in tumoroids cultured in homebrew media (embedded culture). (b) Percentage overlap of single nucleotide variants called in initial tumoroid sample (or starting cell bank) and in tumoroids expanded in OncoPro medium in an embedded culture format. (e) Percentage overlap of single nucleotide variants called in initial tumoroid sample (or starting cell bank) and in tumoroids expanded in OncoPro medium in a suspension culture format.

**a**

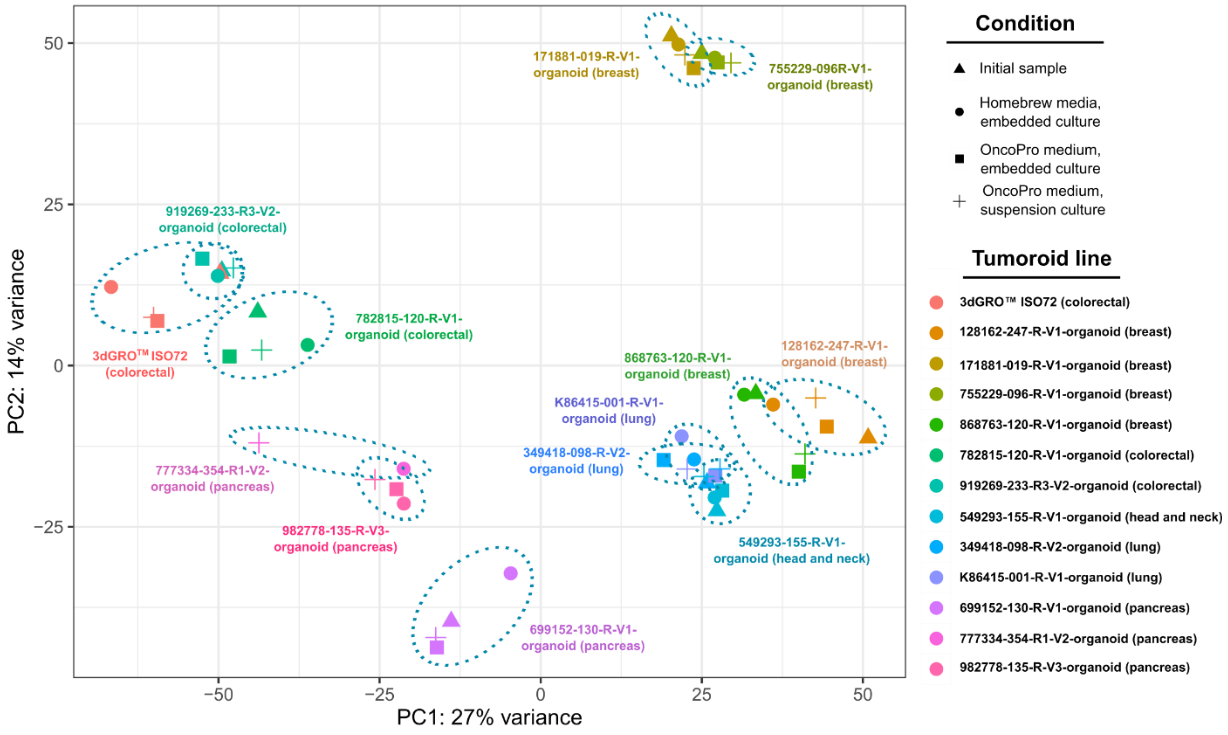


**b**

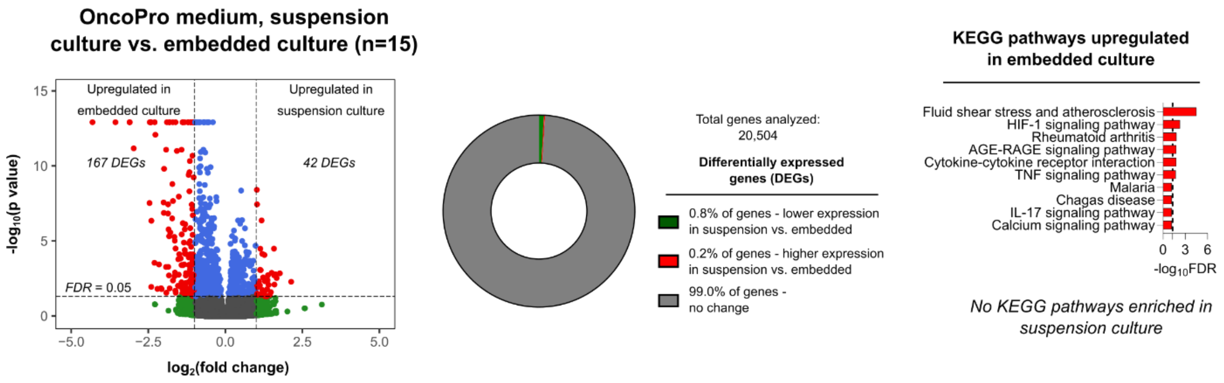


**Supplementary Figure S12. Tumoroids cultured using OncoPro medium retain expected gene expression patterns in suspension and embedded culture formats.** (a) Principal component analysis comparing bulk RNA expression of colorectal (n=3), lung (n=2), breast (n=4), pancreatic (n=3), and head and neck (n=1) tumoroid lines show retention of gene expression profiles irrespective of the culture conditions (suspension and embedded) and culture media (homebrew media and OncoPro medium) in which they were grown. (b) Differential gene expression analysis of tumoroids cultured in OncoPro medium in either suspension or embedded formats across cancer indications (n=15) revealed differentially expressed genes at fold change>2 and false discovery rate (FDR)<0.05. Percentage change in transcriptome and analysis for enriched Kyoto Encyclopedia of Genes and Genomes (KEGG) pathways in highly expressed genes in embedded culture; top 10 results are shown. Dotted line indicates FDR=0.05. No KEGG pathway was enriched when highly expressed genes in suspension culture were used in the analysis.

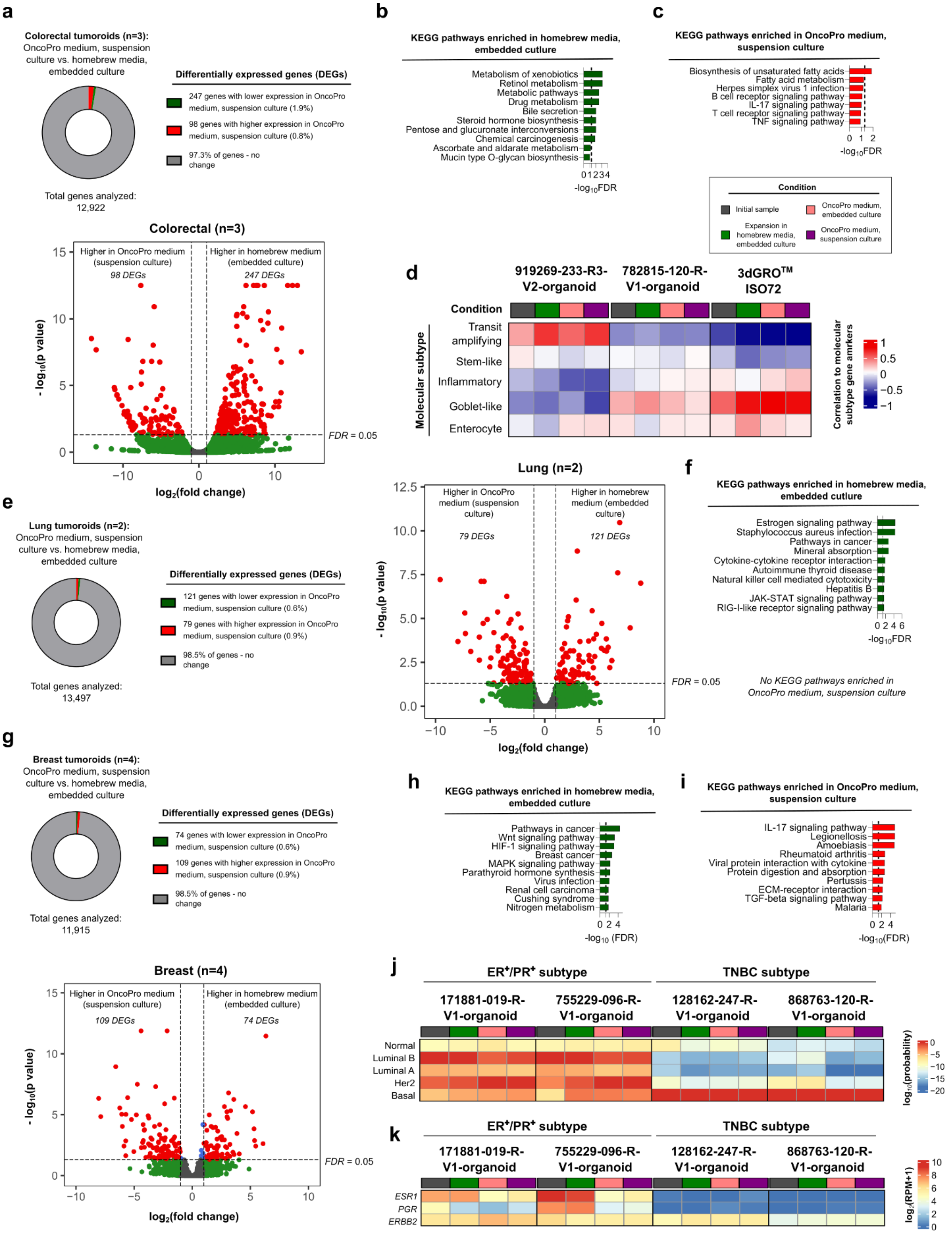

a
Colorectal tumoroids (n=3): OncoPro medium, suspension culture vs. homebrew media, embedded culture
Differentially expressed genes (DEGs)
247 genes with lower expression in OncoPro medium, suspension culture (1.9%)
98 genes with higher expression in OncoPro medium, suspension culture (0.8%)
97.3% of genes - no change
Total genes analyzed: 12,922
Colorectal (n=3)
Higher in OncoPro medium (suspension culture)
98 DEGs
Higher in homebrew medium (embedded culture)
247 DEGs
FDR = 0.05
- log10(p value)
log2(fold change)
b
KEGG pathways enriched in homebrew media, embedded culture
Metabolism of xenobiotics
Retinol metabolism
Metabolic pathways
Drug metabolism
Bile secretion
Steroid hormone biosynthesis
Pentose and glucuronate interconversions
Chemical carcinogenesis
Ascorbate and aldarate metabolism
Mucin type O-glycan biosynthesis
-log10FDR
c
KEGG pathways enriched in OncoPro medium, suspension culture
Biosynthesis of unsaturated fatty acids
Fatty acid metabolism
Herpes simplex virus 1 infection
B cell receptor signaling pathway
IL-17 signaling pathway
T cell receptor signaling pathway
TNF signaling pathway
-log10FDR
Condition
Initial sample
OncoPro medium, embedded culture
Expansion in homebrew media, embedded culture
OncoPro medium, suspension culture
d
919269-233-R3-V2-organoid
782815-120-R-V1-organoid
3dGRO™ ISO72
Condition
Molecular subtype
Transit amplifying
Stem-like
Inflammatory
Goblet-like
Enterocyte
Correlation to molecular subtype gene amrkers
e
Lung tumoroids (n=2): OncoPro medium, suspension culture vs. homebrew media, embedded culture
Differentially expressed genes (DEGs)
121 genes with lower expression in OncoPro medium, suspension culture (0.6%)
79 genes with higher expression in OncoPro medium, suspension culture (0.9%)
98.5% of genes - no change
Total genes analyzed: 13,497
Lung (n=2)
Higher in OncoPro medium (suspension culture)
79 DEGs
Higher in homebrew medium (embedded culture)
121 DEGs
FDR = 0.05
- log10(p value)
log2(fold change)
f
KEGG pathways enriched in homebrew media, embedded culture
Estrogen signaling pathway
Staphylococcus aureus infection
Pathways in cancer
Mineral absorption
Cytokine-cytokine receptor interaction
Autoimmune thyroid disease
Natural killer cell mediated cytotoxicity
Hepatitis B
JAK-STAT signaling pathway
RIG-I-like receptor signaling pathway
-log10FDR
No KEGG pathways enriched in OncoPro medium, suspension culture
g
Breast tumoroids (n=4): OncoPro medium, suspension culture vs. homebrew media, embedded culture
Differentially expressed genes (DEGs)
74 genes with lower expression in OncoPro medium, suspension culture (0.6%)
109 genes with higher expression in OncoPro medium, suspension culture (0.9%)
98.5% of genes - no change
Total genes analyzed: 11,915
Breast (n=4)
Higher in OncoPro medium (suspension culture)
109 DEGs
Higher in homebrew medium (embedded culture)
74 DEGs
FDR = 0.05
- log10(p value)
log2(fold change)
h
KEGG pathways enriched in homebrew media, embedded culture
Pathways in cancer
Wnt signaling pathway
HIF-1 signaling pathway
Breast cancer
MAPK signaling pathway
Parathyroid hormone synthesis
Virus infection
Renal cell carcinoma
Cushing syndrome
Nitrogen metabolism
-log10 (FDR)
i
KEGG pathways enriched in OncoPro medium, suspension culture
IL-17 signaling pathway
Legionellosis
Amoebiasis
Rheumatoid arthritis
Viral protein interaction with cytokine
Protein digestion and absorption
Pertussis
ECM-receptor interaction
TGF-beta signaling pathway
Malaria
-log10(FDR)
j
ER+/PR+ subtype
TNBC subtype
171881-019-R-V1-organoid
755229-096-R-V1-organoid
128162-247-R-V1-organoid
868763-120-R-V1-organoid
Normal
Luminal B
Luminal A
Her2
Basal
log10(probability)
k
ER+/PR+ subtype
TNBC subtype
171881-019-R-V1-organoid
755229-096-R-V1-organoid
128162-247-R-V1-organoid
868763-120-R-V1-organoid
ESR1
PGR
ERBB2
log2(RPM+1)

**Supplementary Figure S13. Differential gene expression analysis following tumoroid expansion in homebrew media or in OncoPro medium suspension culture and characterization by molecular subtype across culture conditions.** (a) Differential gene expression of colorectal tumoroid cultures (n=3) in embedded culture using homebrew media vs. in suspension culture using OncoPro medium revealed differentially expressed genes at fold change>2 and false discovery rate (FDR)<0.05. Percentage change in transcriptome is also displayed. (b) Enriched Kyoto Encyclopedia of Genes and Genomes (KEGG) pathways for colorectal tumoroids in homebrew media, embedded culture. Top 10 results are shown. Dotted line indicates FDR=0.05. (c) Enriched KEGG pathways for colorectal tumoroids in OncoPro medium, suspension culture. Dotted line indicates FDR=0.05. All enriched pathways are shown. (d) Consensus molecular subtypes from gene expression analysis present in cultures of colorectal tumoroids in different culture conditions. (e) Differential gene expression of lung tumoroid cultures (n=2) in embedded culture using homebrew media vs. in suspension culture using OncoPro medium revealed differentially expressed genes at fold change>2 and false discovery rate (FDR)<0.05. Percentage change in transcriptome is also displayed. (f) Enriched KEGG pathways for colorectal tumoroids in homebrew media, embedded culture. Top 10 results are shown. Dotted line indicates FDR=0.05. No KEGG pathway was enriched when highly expressed genes in OncoPro medium suspension culture were used in the analysis. (g) Differential gene expression of breast tumoroid cultures (n=4) in embedded culture using homebrew media vs. in suspension culture using OncoPro medium revealed differentially expressed genes at fold change>2 and false discovery rate (FDR)<0.05. Percentage change in transcriptome is also displayed. (h) Enriched KEGG pathways for breast tumoroids in homebrew media, embedded culture. Top 10 results are shown. Dotted line indicates FDR=0.05. (i) Enriched KEGG pathways for breast tumoroids in OncoPro medium, suspension culture. Top 10 results are shown. Dotted line indicates FDR=0.05. (j) Consensus molecular subtypes from gene expression analysis present in cultures of breast tumoroids in different culture conditions. (k) Expression levels of *ESR1* and *PGR* genes across breast tumoroid lines. Expression of *ERBB2* is displayed for reference.

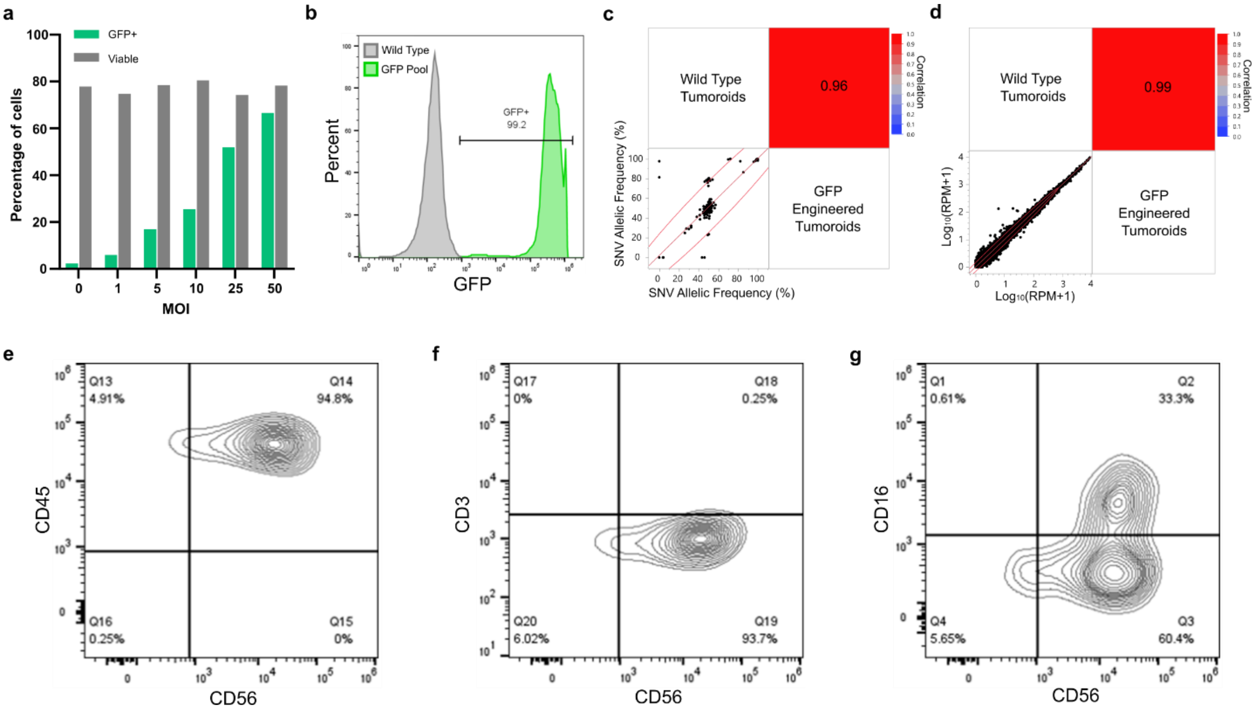


**Supplementary Figure S14. Characterization of tumoroid engineering and primary natural killer (NK) cells.** (**a**) Percentage of viable and green fluorescent protein-positive (GFP+) tumoroid cells 5 days after lentiviral transduction. (**b**) Flow cytometry of wild type or GFP-transduced tumoroids (GFP Pool) after blasticidin selection. (**c**) Correlation of the allelic frequency of single nucleotide variants (SNVs) or (**d**) full transcriptome gene expression levels between wild type or GFP engineered tumoroids (after blasticidin selection). (**e-g**) Flow cytometry characterization of primary NK cells after isolation and expansion.

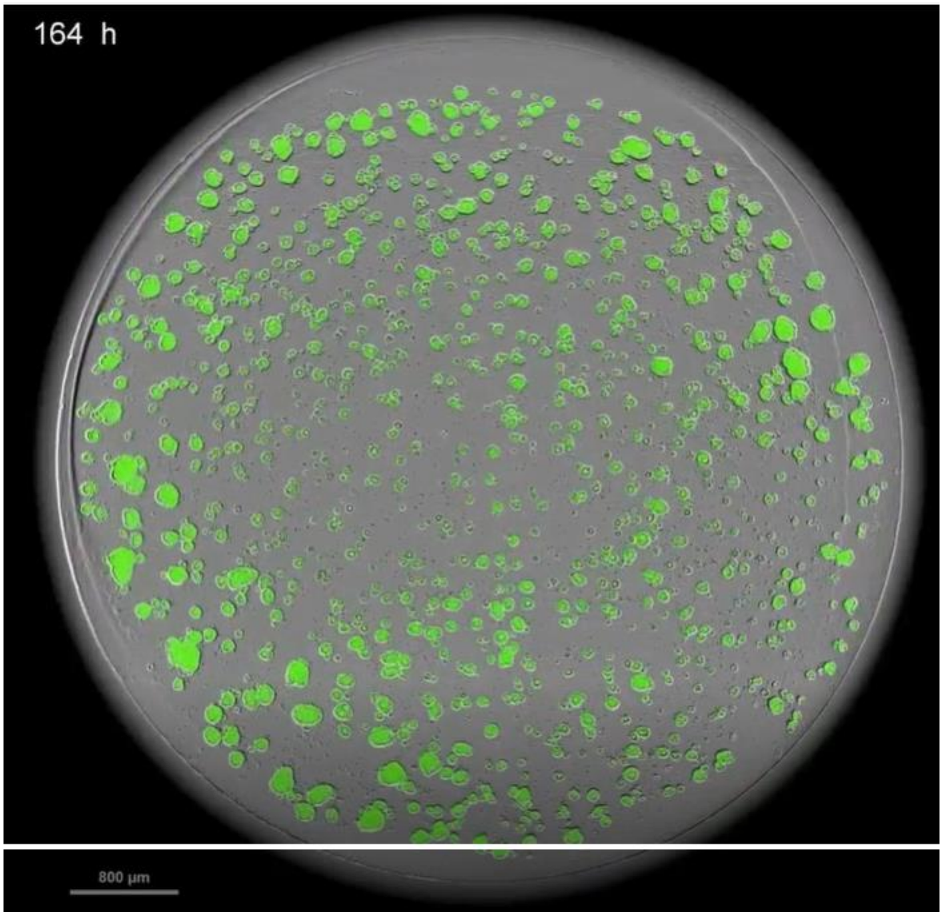


**Supplementary Video S1. Growth of engineered green fluorescent protein-positive (GFP+) tumoroids.** The HuCo1044-GFP tumoroid line was dissociated, seeded into a black walled 96-well plate in suspension culture using OncoPro medium, and imaged every 4 hours. Artifacts in the lower right-hand corner (present during the first 48 hours) are bubbles at the liquid surface. Scale bar = 800 µm.

**Description of Supplementary Tables**

**Supplementary Table S1** – OncoPro medium, indication-specific recommendations

**Supplementary Table S2** – Donor characteristics for tumoroid lines derived in OncoPro medium and described in this manuscript

**Supplementary Table S3** – within- and between-donor correlation values (Pearson's *r*) for variant allele frequency of called single nucleotide variants for OncoPro medium-derived tumoroid lines

**Supplementary Table S4** – Genomic mutations identified in APC gene based on qPCR genotyping assays for select colorectal tumoroids

**Supplementary Table S5** – media recipes used to culture publicly available tumoroids in homebrew conditions

**Supplementary Table S6** – details on culture of NCI PDMR tumoroids and publicly available tumoroid line

**Supplementary Table S7** - within- and between-culture correlation values (Pearson's *r*) for variant allelic frequency of called single nucleotide variants between initial tumoroids (or initial tumoroid working bank), tumoroids expanded in homebrew embedded culture, and tumoroids expanded in OncoPro medium in embedded or suspension formats.
